# NNMT promotes tubular senescence and fibrosis in chronic kidney disease

**DOI:** 10.1101/2025.01.06.631437

**Authors:** Lucie Chanvillard, Hildo Lantermans, Christopher Wall, Jonathan Thevenet, Loes Butter, Loic Tauzin, Nike Claessen, Stefan Christen, Sonia Karaz, Steve Lassueur, Giulia Lizzo, José Luis Sanchez-Garcia, Sylviane Métairon, James A. Holzwarth, Valentina Ferro, Sofia Moco, Erik J.M. van Bommel, Michael J.B. van Baar, Anne C. Hesp, Daniel H. van Raalte, Joris J.T.H. Roelofs, Matthew J. Sanders, Jerome N. Feige, Vincenzo Sorrentino, Alessandra Tammaro

**Author notes:** Correspondence (V.S.), (A.T.). joint supervision of the work (Matthew J. Sanders, Jerome N. Feige, Vincenzo Sorrentino, Alessandra Tammaro).

## Abstract

Chronic kidney disease (CKD) is a major global health issue, projected to become the fifth leading cause of mortality by 2040. Renal tubular cell senescence is a key driver of kidney fibrosis, the final manifestation of CKD. However, current treatment strategies, do not target senescent cells, as the underlying mechanisms driving this dysfunctional phenotype remain poorly described. Here, we identify nicotinamide-N-methyltransferase (NNMT), as a critical mediator of tubular senescence and fibrosis in CKD. Using human RNAseq profiles of CKD, we show that NNMT expression in the renal tubulointerstitium is strongly associated with CKD pathology and transcriptional signatures of cellular senescence. In human diabetic kidney disease biopsies, NNMT levels correlate with the senescence marker p21, kidney function decline, and fibrosis. Spatial transcriptomics further highlights that NNMT-positive tubules are senescent, fibrotic, and surrounded by a pro-inflammatory microenvironment. Preclinical models of early-stage CKD, show upregulation of NNMT and association with senescence. Overexpression of NNMT in TGF-β-stimulated tubular epithelial cells promotes senescence and partial epithelial-to-mesenchymal transition (EMT), while inhibition of NNMT in kidney cells and organoids is protective. Altogether, we identify NNMT as a novel therapeutic target in the early stages of CKD with the potential to reduce tubular senescence, fibrosis and significantly slow disease progression.

## INTRODUCTION

Chronic kidney disease (CKD) is defined as persistent abnormalities in kidney function and/or structure, lasting at least 3 months^1^. CKD is associated with premature aging of other organs and systemic complications, including cardiovascular disease, bone-mineral disorders, and muscle wasting, collectively contributing to the unfavourable prognosis^2^. Approximately 850 million people worldwide are estimated to have kidney disease, roughly double the number of people who live with type 2 diabetes and 20 times more than the prevalence of cancer worldwide, translating to a global prevalence >10%^3^. CKD patients face increased management complexity, experience high comorbidities, longer hospital stays and substantial healthcare costs. It is estimated that in Europe the CKD-related expenditure is surpassing those for cancer or diabetes. Consequently, due to these challenges, CKD has claimed its place on the global public health agenda, highlighting the need for more actions to support early detection and treatment^4^.

Existing therapeutic options aim to delay disease progression rather than targeting the early stages, before kidney failure occurs and replacement therapy via dialysis or kidney transplantation becomes necessary^5^. This underscores the need to investigate molecular mechanisms in the early stages of disease to develop more effective and targeted treatments. Among the different mechanisms contributing to CKD progression, the tubular compartment of the kidney is receiving increasing attention given its active role in renal function and early impact on CKD pathophysiology^6,7^. Renal tubular epithelial cells (TECs) are the most abundant cells in the kidney and control the reabsorption of water, ions and solutes from the urinary filtrate into the bloodstream^8^. In a healthy kidney, TECs have a strong regenerative capacity, allowing them to de-differentiate, proliferate, and re-differentiate in response to stress and injury^9^. However, during the initial stages of CKD, repeated insults to the tubular epithelium, either from increased reabsorption workload or direct injury, lead to a loss of TEC regenerative capacity. As a result, TECs stop cycling and re-differentiating^10^. Chronically injured TECs then enter a state of senescence, characterized by cell cycle arrest and secretion of the senescence-associated secretory phenotype (SASP)^11^. The SASP is rich in pro-inflammatory and pro-fibrotic factors that drive inflammation and immune cell recruitment, and promote a paracrine epithelial-to-mesenchymal transition (EMT) in TECs^12,13^. Tubular senescence and EMT jointly foster parenchymal damage and drive fibrotic remodelling of renal tissue, ultimately leading to kidney functional decline that manifests clinically by lower glomerular filtration rate (GFR) and increased albuminuria^14–17^. While senotherapeutics hold promise for preventing renal fibrotic remodeling and functional impairment by addressing the senescent phenotype^18^, the specific molecular mechanisms driving TEC senescence remain poorly understood. Therefore, identifying novel actionable pathways involved in TEC senescence during early CKD, is crucial for developing new biomarkers and therapies.

The present study reports a novel association between nicotinamide (NAM)-N-methyltransferase (NNMT) and TEC senescence in early-stage CKD. Using human datasets and biopsies, we demonstrate that tubular NNMT expression is associated with clinical and histopathological readouts of CKD, as well as with markers of senescence and fibrosis across multiple kidney disease cohorts. Additionally, we demonstrate a parallel elevation of NNMT and senescence that precedes fibrosis in two preclinical *in vivo* models of early-stage CKD, induced by natural aging and diabetic nephropathy. *In vitro* experiments show that NNMT overexpression in tubular epithelial cells aggravates the senescent and fibrotic response in TECs, while SAM supplementation and NNMT inhibition by a small molecule, significantly reduces senescence and fibrosis in cells. To establish human relevance, the protective effect of NNMTi was further confirmed in a human iPSC-derived kidney organoid model, demonstrating its efficacy in mitigating senescence and EMT, highlighting its therapeutic potential in CKD^19–21^. Our findings mechanistically link tubular NNMT to senescence and fibrosis in kidney disease and rationalize the use of NNMT inhibition as a senotherapeutic intervention to reverse early maladaptive changes in TECs during CKD.

## RESULTS

### NNMT is upregulated in renal tubules of human CKD patients and associates with cellular senescence

In order to identify novel markers or regulators of CKD pathology and cellular senescence, we used publicly available transcriptomic datasets of the kidney tubulointerstitial compartment in individuals diagnosed with CKD via low estimated GFR (eGFR) ^22,23^. Datasets were merged and batch-corrected. A high inter-study clustering in the principal component analysis (PCA) **(Figure 1A)** could be efficiently normalized by a batch correction procedure **(Figure 1B)**. Gene set enrichment analysis (GSEA) of genes ranked by their correlation to eGFR showed a negative enrichment of EMT **(Figure S1A)**, inflammation and cell cycle-related pathways, in line with the expected upregulation of these gene sets with CKD^24^. Conversely, energy metabolism pathways were positively associated with eGFR and thus downregulated in individuals with low renal function. Using gene sets curated for fibrosis and validated for senescence^25–27^, we confirmed a robust activation of these processes in the human tubulointerstitial compartment when kidney function declines **(Figure S1B,C)**. NNMT was the gene that most strongly correlated with eGFR with a very strong negative association, indicating an upregulation in individuals with low kidney function **(Figure 1C,D)**. To understand the disease-specific impacts on this correlation, we calculated the effect of the different covariates on the expression of NNMT **(Figure 1E)**. Besides the previously observed association with eGFR, diabetic kidney disease (DKD), primary etiology of CKD^28^, had the strongest impact on NNMT expression. Additionally, two forms of glomerulonephritis showed a similar but more modest impact on NNMT expression **(Figure 1E)**. To further understand if NNMT expression was directly associated with these dysfunctional pathways, we analyzed gene sets that correlate with NNMT expression and observed a significant enrichment in senescence and fibrosis **(Figure 1F,G)**, suggesting a possible direct link between NNMT and senescence and fibrosis.

**FIGURE 1.**
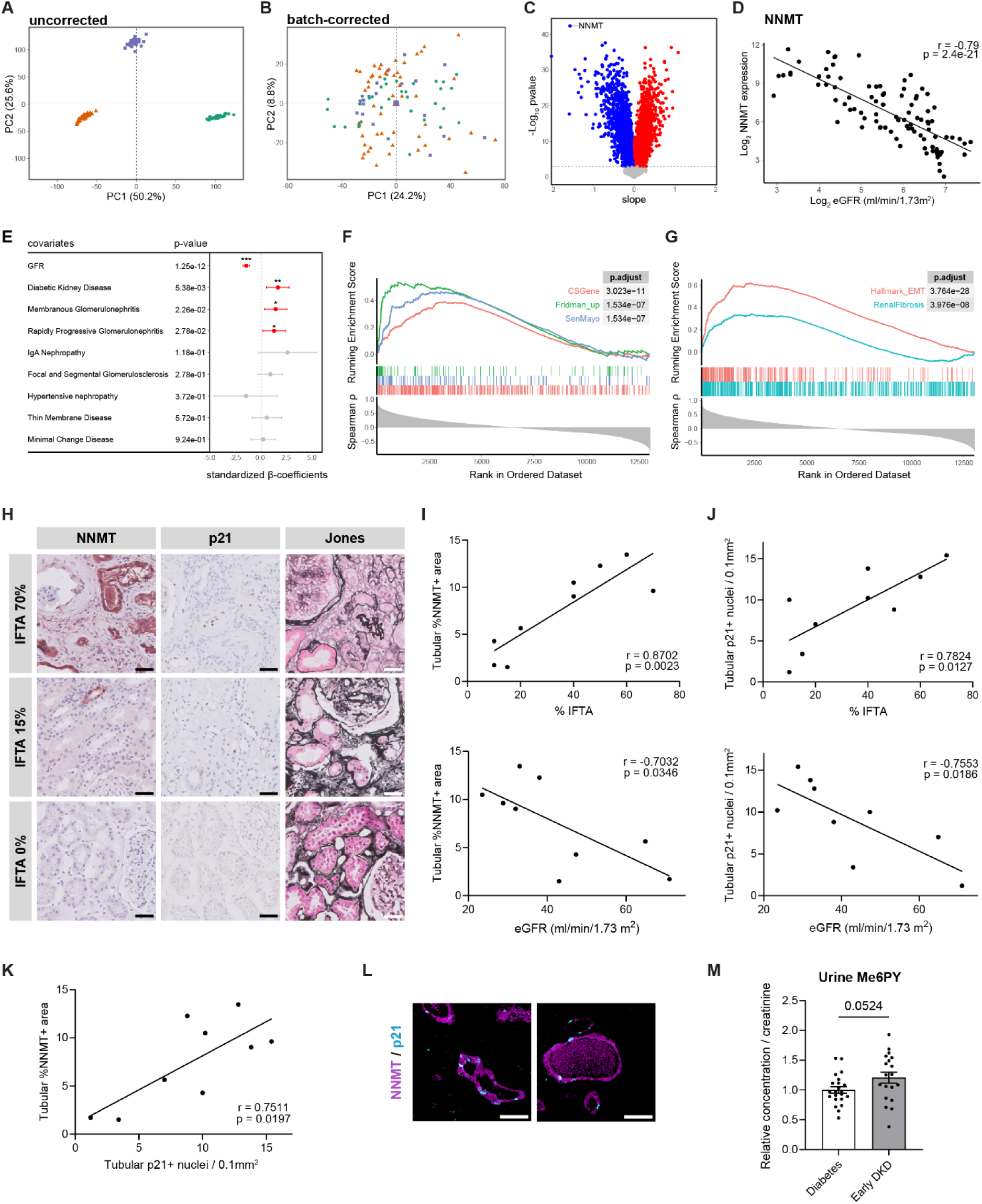
Tubular NNMT associates with human CKD pathology and cellular senescence. **(A,B)** Principal component analysis (PCA) plots (A) before and (B) after batch correction of the three microarray experiments included in the analysis. After batch-adjustment no cluster can be observed. (C) Volcano plot of genes that correlate to eGFR showing slope *vs.* significance of the correlation. n = 104 from the combined dataset. **(D)** Spearman correlation of *NNMT* expression to eGFR. **(E)** Forest plot depicting the effect (β-coefficients with 95% confidence interval) of the disease-specific covariates on the correlation of *NNMT* to eGFR. **(F,G)** GSEA plots, ranking genes by Spearman correlation coefficient to *NNMT* expression and showing enrichment in (F) senescence (CSGene, Fridman_up, SenMayo) and (G) fibrosis (MSigDB epithelial-to-mesenchymal transition hallmark and DisGeNET renal fibrosis) gene sets. **(H)** Representative histological images of DKD kidney biopsies stained for NNMT, p21 and Jones’ stain (used for interstitial fibrosis and tubular atrophy (IFTA) scoring). Scale bar: 50 µm. **(I)** Pearson correlation between tubular % NNMT+ area with % IFTA and with eGFR in human kidney biopsies, n=9. **(J)** Pearson correlation between tubular p21+ nuclei with % IFTA and with eGFR in human kidney biopsies, n=9. **(K)** Pearson correlation of tubular % NNMT+ area with tubular p21+ nuclei in human kidney biopsies, n=9. **(L)** Sequential staining for NNMT and p21 showing colocalization in atrophic tubules. Scale bar: 50 µm. **(M)** LC–HRMS measurement of urinary Me6PY in diabetic (n=22) and early DKD (n=20) patients, mean ± s.e.m., with p-value determined by Student’s two-tailed t-test.

Given the cellular heterogeneity of the kidney, we investigated where NNMT could be upregulated during CKD by analyzing its expression in TECs, interstitial and immune cells using the recently published human kidney cell atlas^29^. The single-cell RNAseq data revealed that the cluster with the highest differential expression of NNMT is the injured proximal tubule (iPT). Within this iPT cluster, NNMT was significantly more expressed in CKD samples compared to control samples **(Figure S1D,E)**. We further validated these findings at the histological level using kidney biopsies from patients with DKD, as this pathology exhibited the most pronounced impact on NNMT expression and represents the leading cause of CKD **(Figure 1E)**. Nine DKD kidney biopsies were analyzed across a wide range of eGFR, and interstitial fibrosis and tubular atrophy (IFTA) scores **(Figure S1F)**. IFTA is considered the main histopathological feature of CKD progression and is therefore incorporated into most pathology scoring systems used to predict renal outcomes, including DKD^30^. IFTA percentage correlated negatively with eGFR in the cohort **(Figure S1G)**. Importantly, the percentage of NNMT-positive area in the tubules significantly correlated with CKD pathology, i.e. positively with IFTA and negatively with eGFR **(Figure 1H,I)**. In addition, we stained for p21, the cyclin-dependent kinase inhibitor responsible for the senescence phenotype in TECs^13,17,31^, and observed increased expression and similar correlations between tubular p21-positive nuclei and CKD pathology **(Figure 1H,J)**. Consistent with our transcriptomic results, NNMT strongly correlated with p21 at the histological level and colocalized at the same atrophic and injured tubules **(Figure 1K,L)**.

NNMT plays a critical role in the metabolism of NAM by catalyzing its methylation, using SAM as a methyl donor, to form 1-methylnicotinamide (MeNAM) **(Figure S1I)**. MeNAM is further oxidized to *N*-methyl-6-pyridone-3-carboxamide (Me6PY; often referred to as Me2PY) in humans (and Me4PY in mice)^32^, which is a uremic toxin that accumulates in the circulation in advanced CKD^33^. To investigate the relevance of the NNMT pathway in the earlier stages of kidney damage (i.e. albuminuria with intact GFR), we measured the metabolites of the NNMT reaction in serum and urine samples from a cohort of patients with type 2 diabetes, either without kidney disease or with early DKD defined by microalbuminuria **(Figure S1H)**. Early DKD patients had higher urinary levels of the terminal metabolite Me6PY compared to diabetic patients **(Figure 1M)**, suggesting that high urinary Me6PY in early DKD could reflect a local rise in NNMT expression within the kidney, before advanced disease stages. Urinary levels of NAM and MeNAM were not affected **(Figure S1J)** and neither NAM, MeNAM nor Me6PY accumulated in the serum of early DKD patients **(Figure S1K)**, possibly because these metabolites may be involved in dynamic metabolic fluxes that do not capture early causes of Me6PY accumulation. Altogether, these data demonstrate that NNMT expression is elevated in injured tubules during CKD in humans, where it associates with declining kidney function, fibrosis and cellular senescence.

### Spatial transcriptomics validates the relationship between NNMT, senescence and the inflammatory microenvironment of injured tubules in human DKD biopsies

To gain deeper molecular insights into NNMT-positive tubules and their surrounding microenvironment, we employed spatial transcriptomics on a subset of the DKD kidney biopsies with a diverse range of fibrosis and NNMT expression markers **(Figure S2A)**. After staining for NNMT, tubular markers (megalin and cytokeratin) and DNA **(Figure 2A)**, regions of interest (ROI) covering either NNMT-positive (NNMT+) or NNMT-negative (NNMT-) tubules were manually selected **(Figure 2B)** and each ROI was automatically segmented into a tubule area of illumination (AOI) and a microenvironment (ME) AOI **(Figure 2C,D)**, and sequenced individually. When comparing the differentially expressed genes in tubular AOIs, we identified a distinct gene expression signature of NNMT+ tubules *vs.* NNMT-tubules **(Figure 2E),** with NNMT being the most significantly differentially expressed gene. Using GSEA on the genes ranked by their fold change in the NNMT+ *vs.* NNMT-AOIs, NNMT+ tubules displayed a significant enrichment of cellular senescence gene sets **(Figure 2C,F)**, validating and confirming the previous findings in an additional cohort of DKD patients **(Figure 1F)** with spatial resolution. Similarly, the most up-regulated pathway in NNMT+ tubules was EMT **(Figure 2G)**, corroborating the association between NNMT and fibrosis **(Figure 1G)**. Interestingly, gene sets related to the transcription factors Myc and TP53 were among the top 10 enriched pathways. Myc is linked to TEC metabolic reprogramming, damage and renal fibrosis^34^, whilst TP53 is known for its contribution to senescence and the EMT program, particularly through p21 activation^35^. Using the Lisa tool^36^, which creates a chromatin landscape model based on the Cistrome Data Browser’s DNase and ChIP-seq database^37^, we found that TP53 is the dominant transcription factor predicted to control gene up-regulation in NNMT+ tubules **(Figure S2B)**. Other gene sets that were upregulated in NNMT+ tubules included those related to interferon gamma and alpha response, TNF-alpha signaling via NF-κB, and the complement system **(Figure 2G),** known to be linked to inflammatory signaling and senescence in kidney disease^38,39^.

**FIGURE 2.**
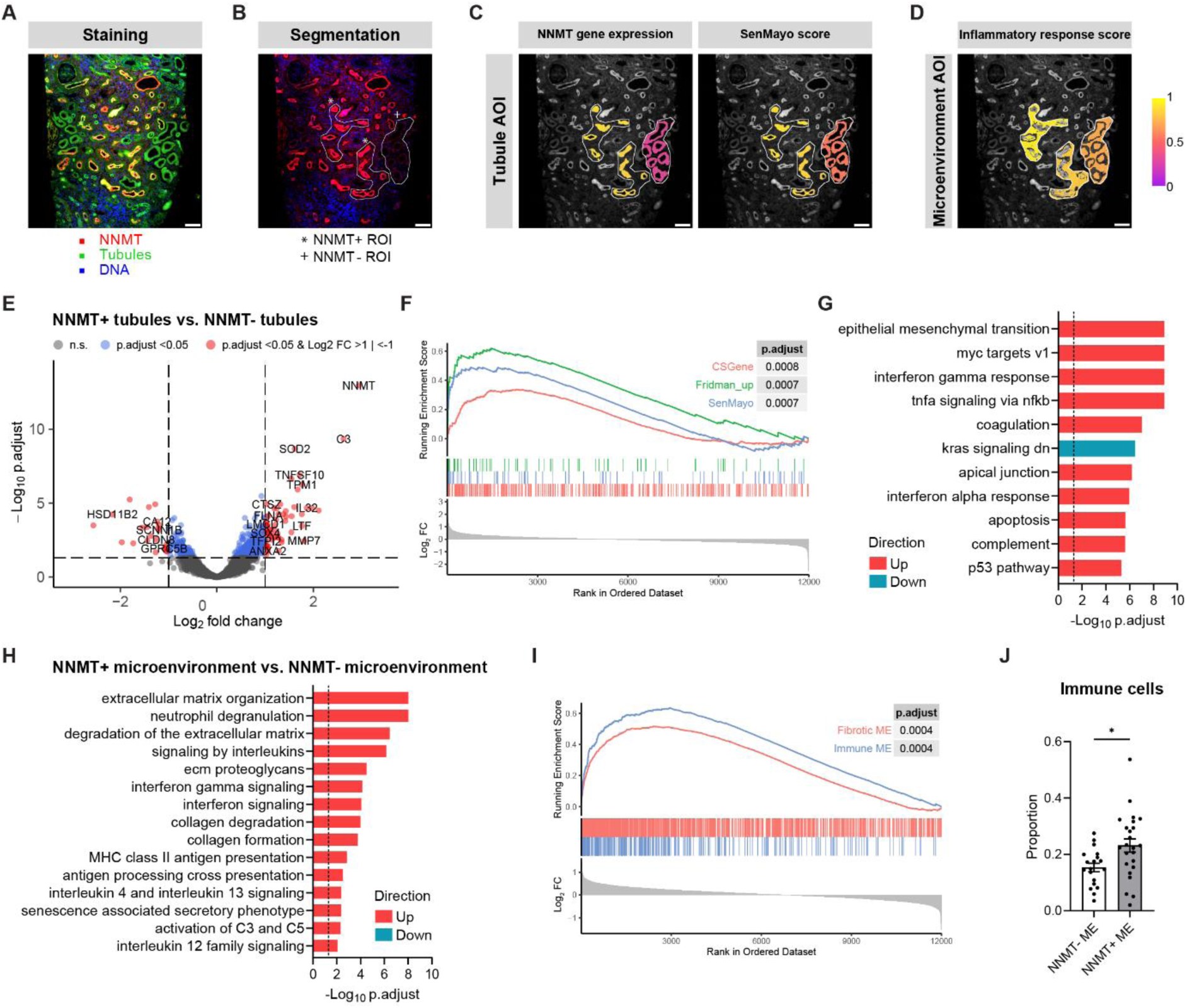
Spatial transcriptomics of human DKD kidney biopsies reveals that NNMT-positive tubules are senescent, fibrotic and surrounded by an inflammatory microenvironment. **(A)** Staining and **(B)** segmentation strategies for spatial transcriptomics analysis of DKD kidney biopsies. Representation of **(C)** the tubule area of illumination (AOI) with a normalized score for *NNMT* gene expression and SenMayo senescence gene set, and **(D)** the microenvironment AOI with an inflammatory response score based on the hallmark gene set from MSigDB. Scale bar: 100 µm. **(E-G)** Transcriptomic analysis of NNMT+ tubules *vs.* NNMT-tubules: (E) volcano plot of differentially expressed genes, (F) GSEA of senescence gene sets (CSGene, Fridman_up, SenMayo), and (G) GSEA of the human hallmark curated gene set collection from MSigDB, showing top gene sets with the lowest p.adjust. **(H,I)** Transcriptomic analysis of NNMT+ microenvironment *vs.* NNMT-microenvironment: (H) GSEA of the human Reactome pathway database showing selected significantly enriched gene sets and (I) GSEA of the fibrotic and immune microenvironment gene sets defined by Abedini *et al.*^29^ **(J)** Computed proportion of immune cells in NNMT+ and NNMT-microenvironment based on the deconvolution analysis, mean ± s.e.m., with p-value determined by Student’s two-tailed t-test.

Given the importance of these pathways in SASP signaling, we hypothesized that senescent NNMT+ tubules may contribute to a sustained inflammatory and fibrotic kidney microenvironment. To investigate this, we examined the differentially expressed genes in the tubular ME AOIs. GSEA of the Reactome collection revealed up-regulation of several pathways linked to inflammation and extracellular matrix remodeling in the NNMT+ ME **(Figure 2D,H)**. When specifically testing for the fibrotic and immune ME gene sets defined by Abedini *et al.*^29^, we confirmed a positive enrichment in the NNMT+ ME compared to the NNMT-ME **(Figure 2I)**. Additionally, cell-type deconvolution of the ME AOIs using a human kidney single-cell RNA-seq dataset^40^ showed a greater proportion of immune cells in the NNMT+ ME compared to the NNMT-ME **(Figure 2J and Figure S2C)**. Finally, to predict the interactions between the tubule and the ME AOIs, we conducted cell-cell communication inference analysis using the CellChat database of receptor-ligand interactions^41^. This analysis identified the strongest interaction signal between the NNMT+ tubules and its ME, specifically within the SASP associated pathways, including collagen, complement, and chemokines and cytokines (CXCL, CCL) networks **(Figure S2D)**.

Collectively, our spatial transcriptomic analyses on DKD kidney biopsies demonstrate that tubules expressing NNMT are more senescent and fibrotic, and that their microenvironment is enriched in inflammatory and immune mediators.

### Renal NNMT and senescence are increased in preclinical models of early-stage CKD

Following our observation of increased urinary Me6PY in early DKD patients **(Figure 1M)**, we asked whether NNMT expression in the kidney is a marker for the initial stages of senescence-associated kidney function decline using two preclinical models of early CKD: i) 16-week-old genetically diabetic (*db/db*) mice^42^ with hyperfiltration and moderate albuminuria **(Figure 3A,B)**, but no increase in renal collagen deposition **(Figure 3E)**, and ii) 24-month-old WT mice^43^ with no detectable kidney function impairment based on creatinine and albuminuria **(Figure 3C,D)** but early fibrotic kidney lesions detected by picrosirius red **(Figure 3F)**. Despite their phenotypic differences, at the molecular level both models had increased renal senescence as evidenced by transcriptomic enrichment of senescence gene sets **(Figure S3A,B)** and increased p21 protein levels **(Figure 3G,I and Figure S3C,D)**. NNMT protein expression were also elevated in *db/db* and aged mice compared to control *db/m* and young mice, respectively **(Figure 3G,I and Figure S3C,D)**. Consistently, the levels of NNMT-dependent metabolites shifted in the kidney of both models, with increased levels of the downstream MeNAM substrate and reduced levels of the upstream methyl donor SAM, while NAM and NAD^+^ levels appeared unchanged **(Figure S3E,F)**. Finally, in both *db/db* and aged mice kidney, we observed a strong correlation between NNMT and p21 protein levels **(Figure 3H,J)**. Using whole genome transcriptomics, senescence gene sets were also enriched in genes significantly correlated with NNMT expression in these early preclinical models of renal decline **(Figure 3K,L)**, consistent with what we had observed in human CKD datasets **(Figure 1F)**.

**FIGURE 3.**
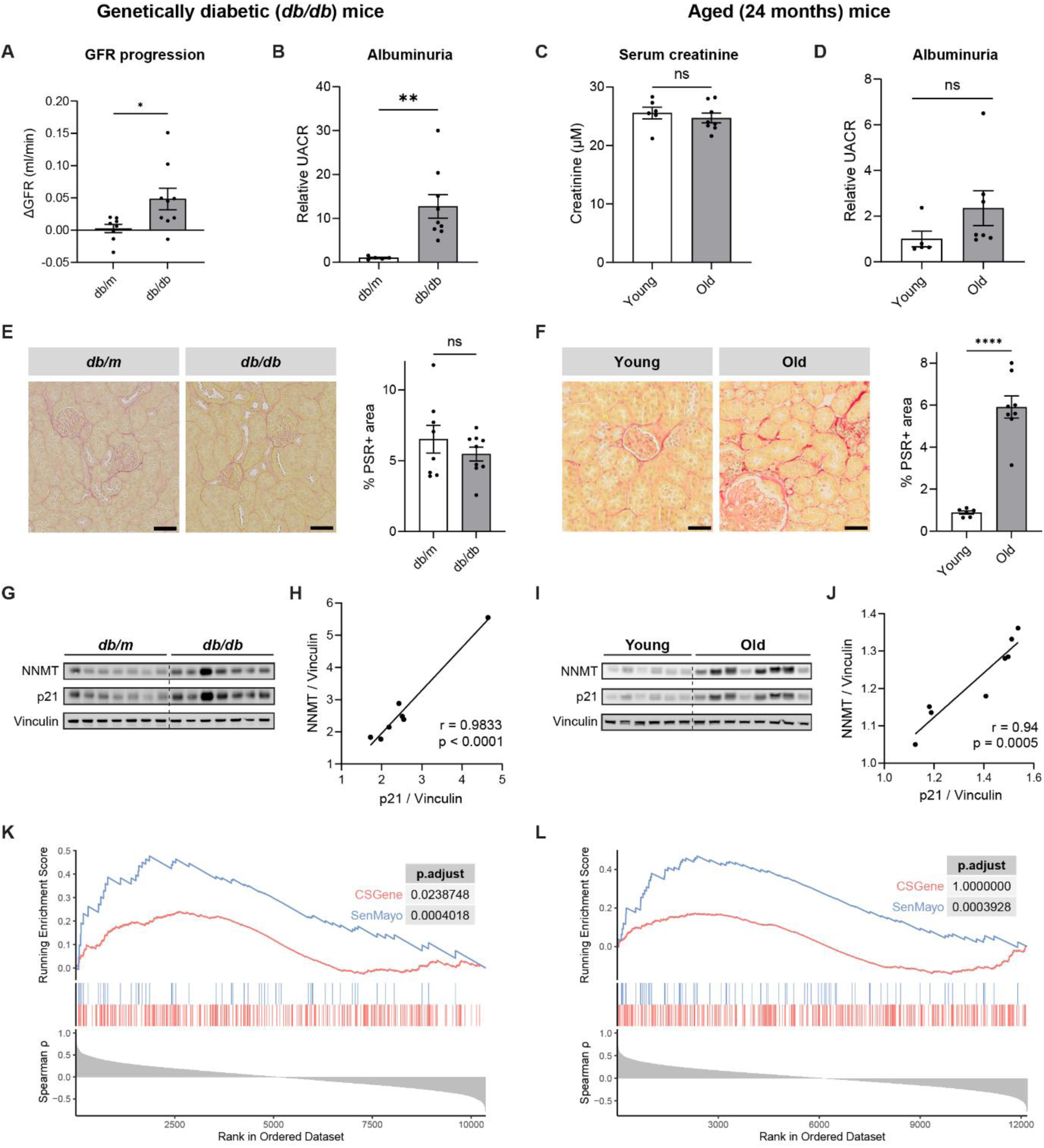
NNMT and senescence correlate in kidneys from diabetic (*db/db*) and aged (24 months) mice. **(A,B)** Kidney function metrics in *db/db* mice and control (*db/m*) mice: (A) GFR progression between mid and end study and (B) urinary albumin/creatinine ratio at the endpoint of the study, n=5-9. **(C,D)** Kidney function metrics in aged (24 months) and young (7 months) mice: (C) serum creatinine and (D) urinary albumin/creatinine ratio at the endpoint of the study, n=5-8. **(E)** Picro-Sirius red (PSR) staining and quantification of collagen deposition based on PSR+ area in *db/m* and *db/db* mice kidney, scale bar: 50 µm, n=8-9. **(F)** PSR staining and quantification of PSR+ area in young and aged mice kidney, scale bar: 50 µm, n=6-8. **(G)** Western blot analysis of NNMT, p21 and vinculin in *db/m* and *db/db* mice. (**H)** Pearson correlation of the normalized expression of NNMT and p21 in *db/db* mice, n=7. **(I)** Western blot analysis of NNMT, p21 and vinculin in young and aged mice. **(J)** Pearson correlation of the normalized expression of NNMT and p21 in aged mice, n=6-8. **(K,L)** GSEA plots, ranking genes by Spearman correlation coefficient to *Nnmt* expression and showing enrichment in senescence (CSGene, SenMayo) gene sets in (K) *db/db* mice kidney and (L) aged mice kidney. Results shown are mean ± s.e.m., with p-value determined by Welch’s two-tailed t-test (A,B,F) or Student’s two-tailed t-test (C-E).

Taken together, these data demonstrate that increased renal NNMT is a biomarker of impaired kidney function and fibrosis, and directly associates with cellular senescence across preclinical models of early CKD and across species.

### NNMT overexpression exacerbates the senescent and pro-fibrotic phenotype in tubular epithelial cells

To investigate the causal relationship between NNMT, senescence and fibrosis, we challenged TECs with a low dose of TGFβ, a model that mimics the senescent and pro-fibrotic phenotypic shift in TECs^44,45^. Our *in vitro* model recapitulated the transcriptomic changes observed in tubules during renal decline with increased senescence, EMT and inflammation, decreased expression of metabolic pathways **(Figure S4A-C)**, and elevation of NNMT levels **(Figure S4D,E)**.

To assess if increased NNMT expression is sufficient to drive senescence and fibrosis, we overexpressed *Nnmt* genetically (*Nnmt*-OE) via lentiviral transduction with a tagged *Nnmt* cDNA. Overexpression of *Nnmt* was confirmed by a 50-fold increase in NNMT protein levels, detection of the myc-tag, and a drastic reduction in the ratio of NAM / MeNAM and SAM / S-adenosyl-homocysteine (SAH) metabolites **(Figure 4A-C)**. Given the critical role of NAM in the generation of NAD^+^ via the nicotinamide phosphoribosyltransferase (NAMPT)-mediated salvage pathway in mammals^46^, we also measured NAD^+^ in this model. However, no changes were detected **(Figure 4D)**, consistent with observations *in vivo* in the kidney of *db/db* and aged mice **(Figure S3E,F)**. At the transcriptomic level, *Nnmt*-OE cells in absence of TGFβ had a small but significant enrichment in senescence and fibrosis gene sets **(Figure 4E)**, and phenotypically, a decline in cell proliferation determined by an EdU pulse **(Figure 4G)**. However, when stimulated for 48 hours with TGFβ, *Nnmt*-OE aggravated the senescent and dedifferentiated phenotype. Indeed, in addition to the transcriptomic enrichment of curated fibrosis and senescence genesets **(Figure 4F)**, we observed a further decrease in cell proliferation, an increase in alpha smooth muscle actin (αSMA) protein levels (a marker of EMT), elevated p21 protein levels, and enhanced senescence-associated β-galactosidase (SA-β-gal) activity, compared to mock cells treated with TGFβ **(Figure 4G-I)**.

**FIGURE 4.**
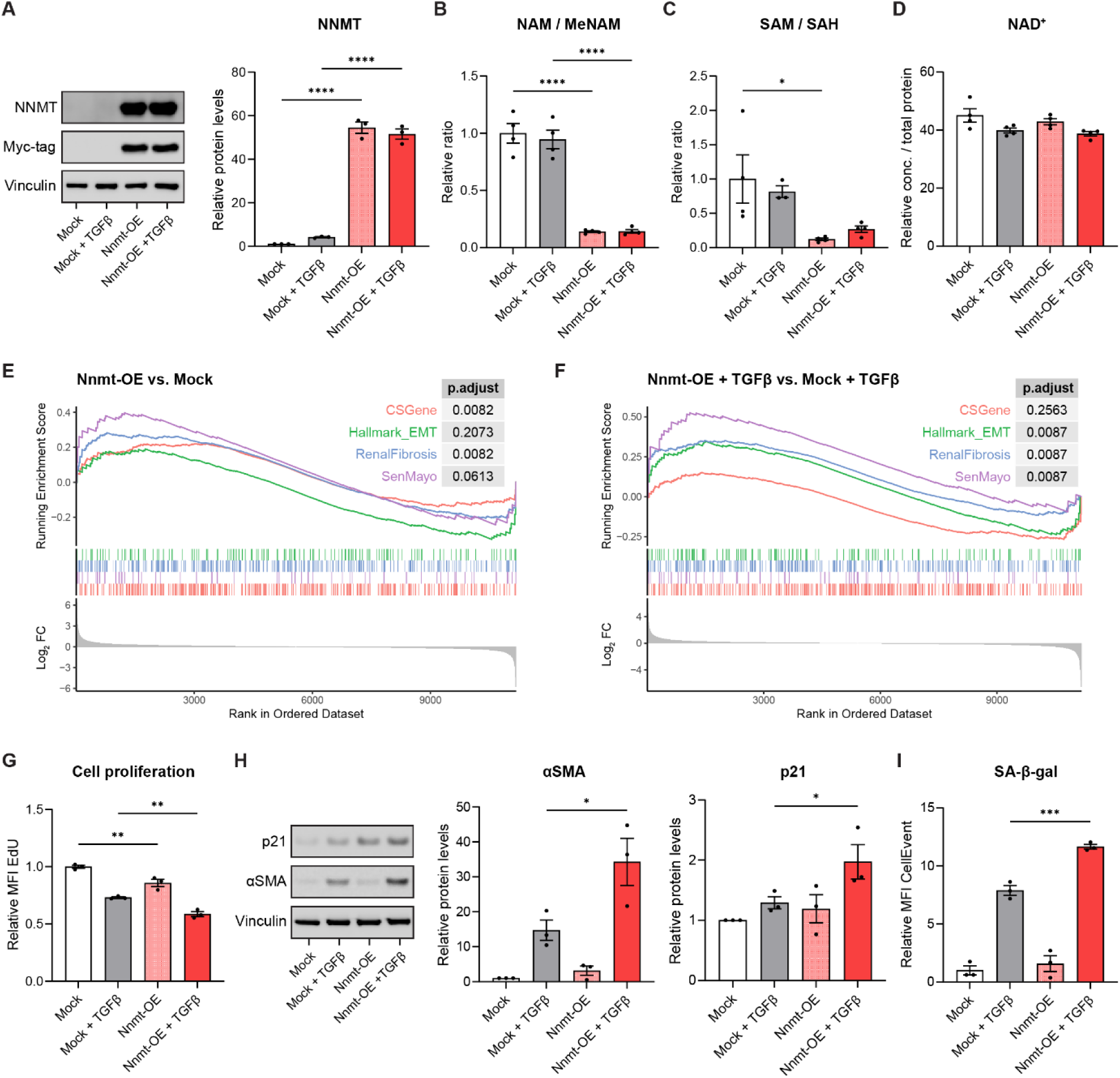
NNMT overexpression aggravates senescence and EMT in tubular epithelial cells. **(A)** Western blot analysis of NNMT, myc-tag and vinculin in *Nnmt* overexpressing (*Nnmt*-OE) and mock-transfected tubular epithelial cells, with and without TGFβ stimulation, and quantification of NNMT protein levels, n=3. **(B-D)** LC–HRMS measurement of (B) NAM / MeNAM ratio, (C) SAM / SAH ratio and (D) NAD^+^ in cells as in A, n=4. **(E,F)** GSEA of senescence (CSGene, SenMayo) and fibrosis (Hallmark_EMT, RenalFibrosis) gene sets in (E) *Nnmt*-OE *vs.* mock cells and (F) *Nnmt*-OE *vs.* mock cells, both stimulated with TGFβ, n=3. **(G)** Flow cytometry analysis of EdU in the same conditions as in A, n=3. **(H)** Western blot analysis of αSMA, p21 and vinculin and quantification of protein levels in cells as in A, n=3. **(I)** Flow cytometry analysis of CellEvent in the same conditions as in A, n=3. Results shown are mean ± s.e.m., with p-value determined by one-way ANOVA with Šídák’s post-hoc test (A-D,G-I).

Overall, these *in vitro* results demonstrate that NNMT over-expression is sufficient to prime TECs to pro-fibrotic and senescence signaling by modulating SAM and MeNAM levels, and this is aggravated to functional impairments via TGFb signaling.

### The NNMT substrates NAM and methyl-donor SAM have diverse outcomes on senescence and fibrosis

Given that the NNMT-mediated enzymatic reaction consumes NAM and SAM and the changes of these metabolites observed both *in vivo* **(Figure S3E,F)** and in our *in vitro Nnmt*-OE model **(Figure 4B,C)**, we tested if exogenous supplementation of these substrates could counteract high NNMT and support tubular homeostasis. 21-month aged mice supplemented with the NNMT substrate NAM in the diet for 14 weeks were analyzed using histological, molecular and metabolic readouts **(Figure 5A)**. NAM levels did not change in the kidney of NAM-treated mice, but both MeNAM and NAD^+^ levels were increased **(Figure S5A)**, confirming the bioavailability of NAM in the kidney, with a higher flux through both NNMT and NAMPT^47,48^. In agreement with what observed in our first aging study (**Figure 3F)**, aged kidneys analyzed by histology had interstitial fibrosis and tubular atrophy (IFTA), but these phenotypes were not ameliorated by NAM **(Figure 5B,C)**. Furthermore, NAM supplementation did not affect senescence and fibrosis gene sets at the transcriptomic level **(Figure 5D)**. Quantification of protein expression by western blot confirmed that αSMA levels were not changed in the kidney of aged mice supplemented with NAM **(Figure S5B,C)**. Consistently, the same amount of p21+ nuclei was observed in the renal tubulointerstitium of aged and NAM-supplemented aged mice **(Figure 5E)**. These results therefore demonstrate that NAM supplementation does not rescue the defects of NNMT over-expression in early CKD and suggest that depletion of NAM does not drive senescence and fibrosis phenotypes when NNMT is over-expressed.

**FIGURE 5.**
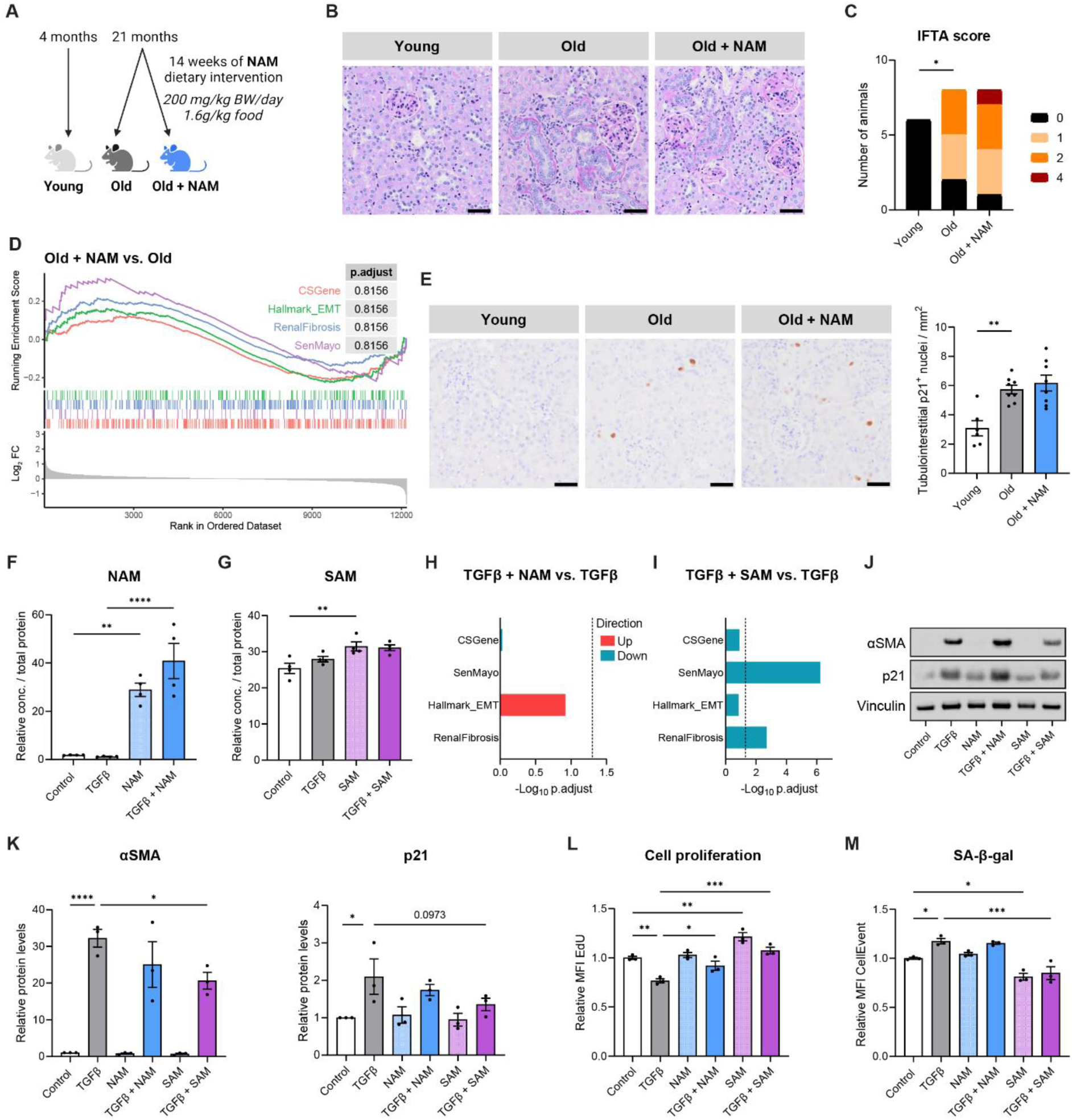
NAM supplementation does not protect against renal fibrosis and senescence while SAM mitigates the damage phenotype. **(A)** Experimental design of the preclinical study of dietary NAM intervention in aged mice, n=6 (young), n=8 (old, old+NAM). **(B)** Representative images of kidney histological assessment based on PAS-D staining in the kidney of young mice, old mice, and old mice supplemented with NAM, scale bar: 50 µm. **(C)** Interstitial fibrosis and tubular atrophy (IFTA) score of the kidney of mice as in B, determined by the pathologist (J.J.T.H.R.) in a blinded fashion. Score is based on a scale of 1 to 4 after PAS-D staining. **(D**) Senescence (CSGene, SenMayo) and fibrosis (Hallmark_EMT, RenalFibrosis) GSEA plots of old mice treated with NAM *vs.* old mice. **(E)** Representative images of p21 immunostaining in the kidney of mice as in B, scale bar: 50 µm, and quantification of tubulointerstitial p21+ nuclei. **(F)** LC–HRMS measurement of NAM in tubular epithelial cells after 48-h TGFβ treatment, with or without co-treatment with NAM, n=4. **(G)** LC–HRMS measurement of SAM in cells after 48-h TGFβ treatment, with or without co-treatment with SAM, n=4. **(H,I)** Senescence and fibrosis GSEA in (H) TGFβ + NAM *vs.* TGFβ treated cells, and (I) TGFβ + SAM *vs.* TGFβ treated cells. **(J)** Western blot analysis of αSMA, p21 and vinculin and **(K)** quantification of protein levels in cells after 48-h TGFβ treatment, with or without co-treatment with NAM or SAM, n=3. **(L,M)** Flow cytometry analysis of (L) EdU and (M) CellEvent in the same conditions as in K, n=3. Results shown are mean ± s.e.m., with p-value determined by one-way ANOVA with Šídák’s post-hoc test (E-G,K-M) or by Kruskal-Wallis (nonparametric) with Dunn’s post-hoc test (C). For the *in vitro* readouts, multiple comparisons were performed between the control group and the TGFβ, NAM and SAM groups, as well as between the TGFβ group and the TGFβ+NAM and TGFβ+SAM groups.

We then assessed if NAM or SAM could prevent senescence and EMT in TECs treated with TGFβ. NAM and SAM were successfully taken up by TECs and increased the levels of these metabolites **(Figure 5F,G)**. In agreement with the *in vivo* results, NAM supplementation in TGFβ-treated cells increased both MeNAM and NAD^+^ levels **(Figure S5D)**, but failed to reverse most phenotypes induced by TGFβ in TEC cells **(Figure 5H,J-M)**. Conversely, SAM significantly reversed the downregulation of senescence and renal fibrosis gene sets in TGFb-treated TEC cells **(Figure 5I)**, which translated functionally into reduced αSMA levels and a similar trend for p21 **(Figure 5J,K)**. In addition, proliferation of cells co-treated with TGFβ and SAM, was restored to healthy cell rates **(Figure 5L)**, and SA-β-Gal activity was significantly reduced **(Figure 5M)**.

In summary, these supplementation studies demonstrate that NAM and the resulting increase in NAD^+^ levels are not sufficient to prevent tubular senescence and fibrotic phenotypes while SAM administration can successfully attenuate this damaging phenotype.

### NNMT inhibition protects against senescence and EMT

Since SAM has a short half-life and would be difficult to use therapeutically, we then used a more targeted approach using a small-molecule inhibitor of NNMT activity to test if targeting the upregulation NNMT in CKD would be beneficial. The selective and membrane-permeable 5-amino-1-methylquinolinium iodide NNMT inhibitor (NNMTi)^19,20^ was used at a non-toxic dose of 50 µM **(Figure S6A)**. As expected, NNMT inhibition in the TGFβ-induced senescence significantly increased the NAM / MeNAM ratio **(Figure 6A)**, and elevated the SAM / SAH ratio **(Figure 6B)**. RNAseq analyses revealed that NNMTi down-regulated a broad set of genes in cells co-treated with TGFβ **(Figure 6C)**. Consistently, these changes were linked to down-regulation of inflammation, EMT and senescence pathways by GSEA **(Figure 6D,E)**. These transcriptional signatures translated into reduced expression of tubular EMT markers, αSMA, Twist and Snail at the protein level, together with a decrease in p21 in TGF-b stimulated TECs co-treated with NNMTi **(Figure 6F)**. Notably, cell proliferation increased by 70% in cells co-treated with TGFβ and NNMTi, and SA-β-Gal activity was restored to healthy control levels **(Figure 6G,H)**.

**FIGURE 6.**
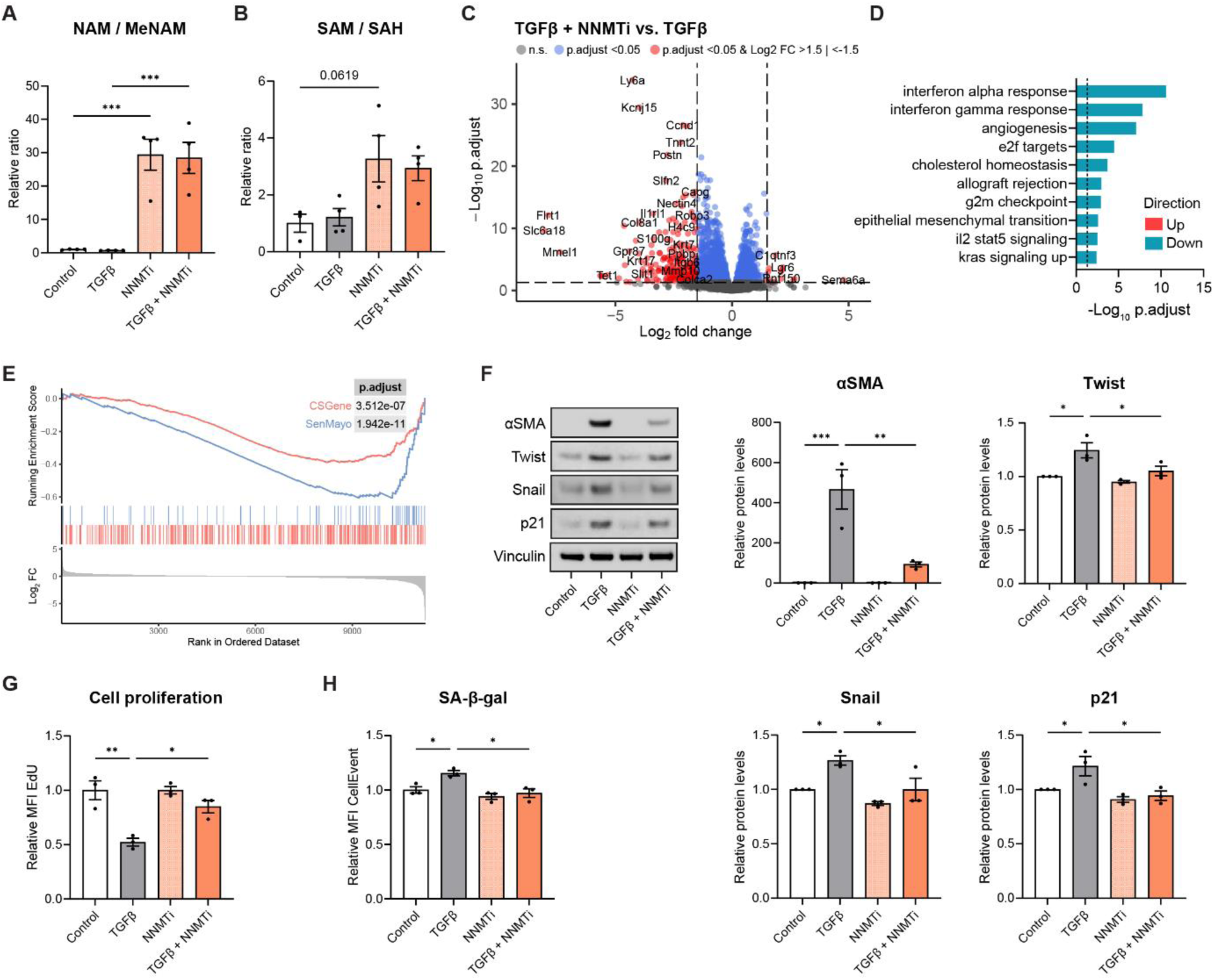
NNMT inhibition protects against senescence and EMT in TGFβ-stimulated tubular epithelial cells. **(A,B)** LC–HRMS measurement of (A) NAM / MeNAM ratio and (B) SAM / SAH ratio in control tubular epithelial cells, cells treated with TGFβ, NNMTi, or both, n=4. **(C-E)** Transcriptomic analysis of TGFβ and NNMTi-treated cells *vs.* TGFβ-treated cells, n=3: (C) volcano plot of differentially expressed genes, (D) GSEA of the mouse hallmark curated gene set collection from MSigDB, showing top 10 gene sets with the lowest p.adjust, and (E) GSEA of senescence gene sets (CSGene, SenMayo). **(F)** Western blot analysis of EMT markers (αSMA, Twist, Snail), senescence (p21) and vinculin and quantification of protein levels in cells as in A, n=3. **(G,H)** Flow cytometry analysis of (G) EdU and (H) CellEvent in the same conditions as in A, n=3. Results shown are mean ± s.e.m., with p-value determined by one-way ANOVA with Šídák’s post-hoc test (A,B,F-H).

Following our observation that NNMT overexpression did not alter NAD^+^ levels (**Figure 4D**), we reasoned that NNMT inhibition should not affect the NAD^+^ pool and mitochondrial metabolism. As expected, NNMTi did not affect NAD^+^ levels in TGFβ-treated cells **(Figure S6B)**. Similarly, mitochondrial respiration measured by Seahorse respirometry was unchanged in the NNMTi co-treated cells **(Figure S6C)**. GSEA also did not detect any enrichment of mitochondrial pathways such as the fatty acid metabolism and oxidative phosphorylation hallmark gene sets **(Figure S6D)**, nor any enrichment of the NAD^+^ metabolic process in NNMTi co-treated cells **(Figure S6E)**. Lastly, while TGFβ stimulation significantly reduced carnitine and acylcarnitine levels, key fatty acid oxidation intermediates^49^, NNMTi did not further alter these metabolite levels **(Figure S6F)**.

To further evaluate the effect of NNMTi in a more complex and human-relevant system, we used iPSC-derived kidney organoids, which have been shown to effectively recapitulate the initiation and progression of renal fibrosis^50,51^. We established a model of senescence and EMT by subjecting the organoids to hypoxic conditions (1% O2) for 48 hours, followed by 24 hours of reoxygenation. Hypoxia is a key factor in CKD pathogenesis, driven by reduced oxygen availability from peritubular capillary loss and tubulointerstitial inflammation, as well as increased oxygen demand to sustain TEC workload and manage oxidative stress^52^. During the reoxygenation phase following hypoxic stress, injured TECs undergo senescence and EMT, providing a model to test NNMT modulation as a senotherapeutic approach **(Figure 7A)**.

**FIGURE 7.**
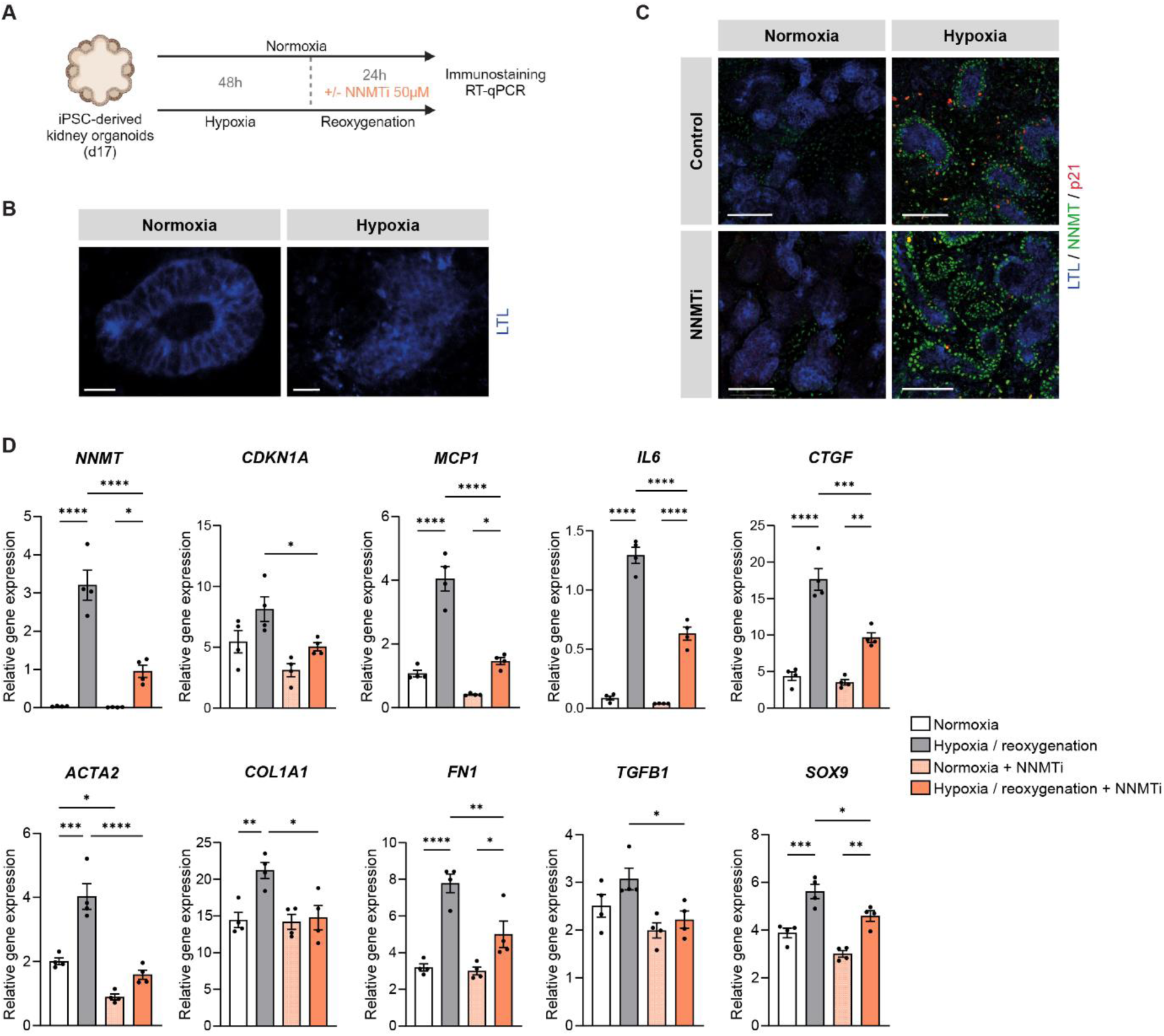
NNMT inhibition protects against senescence and fibrosis after hypoxia/reoxygenation damage in human kidney organoids. **(A)** Experimental setup: human kidney organoids are subjected to 48 hours of hypoxia (1% O2), followed by supplementation with NNMTi during the subsequent 24 hours of reoxygenation. Normoxic controls are kept in standard culture conditions and supplemented with NNMTi 24h before the experimental readouts. **(B)** Representative images of an LTL-stained proximal tubule in kidney organoids under normoxia or hypoxia/reoxygenation conditions, scale bar: 20 µm, magnification x60. **(C)** LTL, NNMT and p21 whole-mount staining in kidney organoids under normoxia or hypoxia/reoxygenation conditions with and without NNMTi, scale bar: 100 µm, magnification x60. **(D)** mRNA expression of *NNMT*, and senescence, SASP and fibrosis genes, *CDKN1A*, *MCP1*, *IL6*, *CTGF*, *ACTA2*, *COL1A1*, *FN1*, *TGFΒ1*, and *SOX9* in kidney organoids under normoxia or hypoxia/reoxygenation conditions with and without NNMTi. n=4 organoids. Results shown are mean ± s.e.m., with p-value determined by one-way ANOVA with Šídák’s post-hoc test (D).

Proximal tubule damage in the organoids following hypoxia/reoxygenation was demonstrated by lotus tetragonolobus lectin (LTL) immunostaining, which showed a loss of cellular integrity and brush border structure **(Figure 7B)**. Representative whole-mount images of the organoids revealed overall increase of NNMT and tubular p21 protein levels under hypoxia/reoxygenation conditions compared to normoxic controls, while NNMTi treatment reduced p21 expression without impacting NNMT protein levels **(Figure 7C)**. These findings were further validated by RT-qPCR, which showed elevated transcript levels of *NNMT* and genes associated with senescence (*CDKN1A*: p21), pro-inflammatory SASP (*MCP1*, *IL6)*, pro-fibrotic SASP (*CTGF, TGFΒ1*) and EMT / fibrosis (*ACTA2*: αSMA, *COL1A1*, *FN1)* in kidney organoids exposed to hypoxia/reoxygenation and reversal by NNMT inhibition **(Figure 7D)**. A similar pattern was observed for *SOX9*, recently identified as a key marker of injured TECs driving renal fibrotic remodeling^53^ **(Figure 7D)**. Consistent with what was observed in TECs, NNMTi treatment did not affect NAD^+^ levels in kidney organoids **(Figure S6G)**.

Taken together, our results demonstrate that NNMT inhibition provides significant protection against senescence and EMT, effectively restoring tubular homeostasis in models of damaged TECs and kidney organoids recapitulating hallmarks of human CKD.

## DISCUSSION

CKD is projected to become the fifth leading cause of mortality by 2040, making its early detection and effective treatment a global public health priority^4,54^. Given its instrumental role in kidney function decline, tubular homeostasis has become the focal point of kidney pathology research to support therapeutic approaches development^7,55^. Cellular senescence, a key pathological phenotype affecting TECs, and is closely studied for its early role in driving fibrotic remodeling and functional decline, highlighting its potential as a target for novel treatments^18,56^.

In this study, we identified NNMT as a tubular marker strongly associated with kidney pathology and cellular senescence across multiple cohorts of human CKD. NNMT expression has been previously found to be elevated in several cancers and associated with poor prognosis^57^, and several recent studies have linked NNMT expression to renal fibrosis^58–61^. However, its role and implications in chronic diseases remain poorly understood^62^. Here, we provide novel evidence that NNMT is associated with tubular senescence and EMT, essential factors in CKD progression (**Figure 8**), and that its inhibition represents a promising senotherapeutic strategy to mitigate this detrimental cellular phenotype and its negative consequences of CKD physiopathology.

**FIGURE 8.**
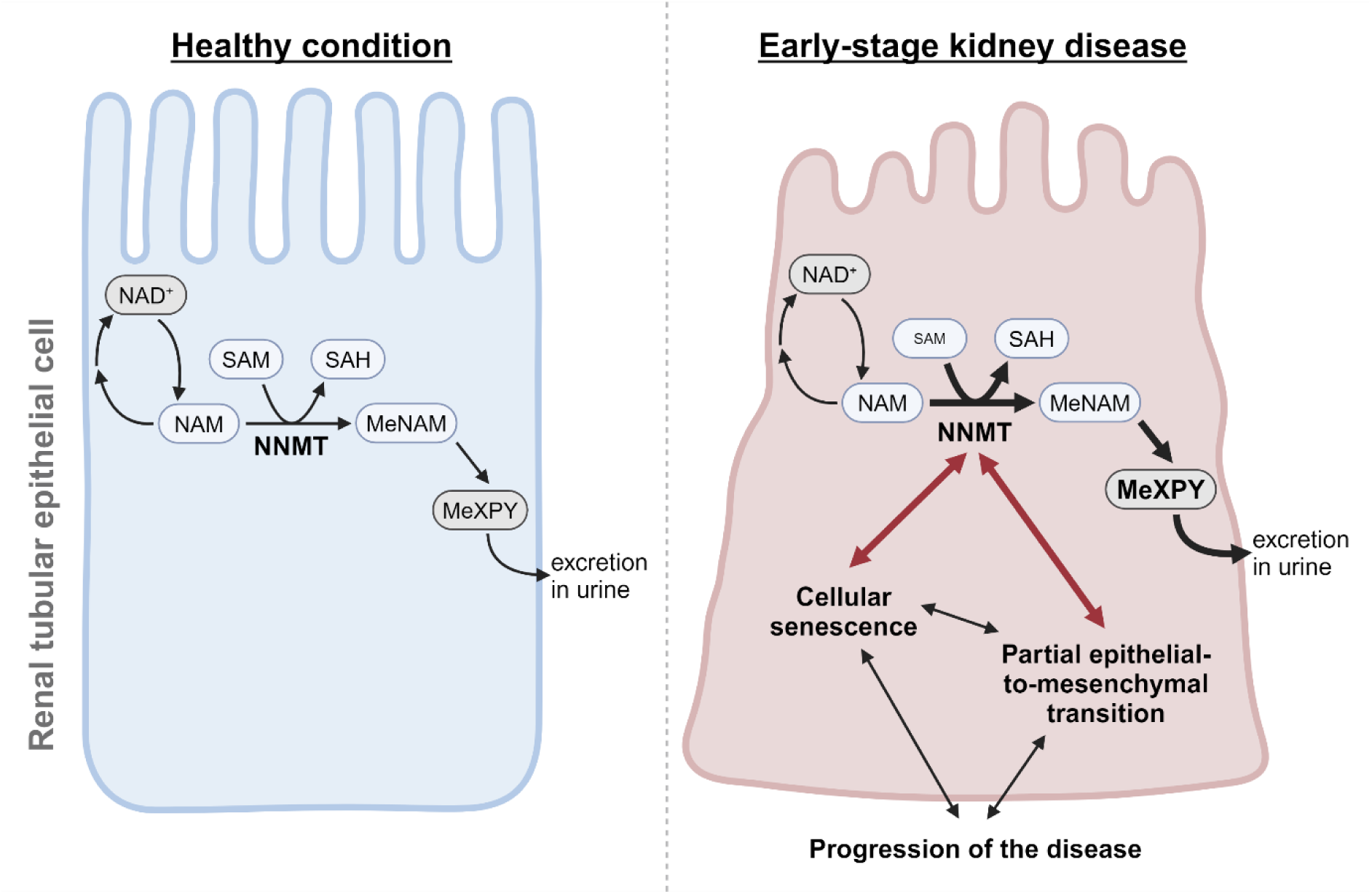
Model summarizing the alterations of kidney tubular epithelial cell homeostasis in response to increased NNMT expression in early-stage kidney disease. Increased flux in the NNMT-catalyzed reaction leads to increased levels of MeXPY in the urine, and decreased intratissular SAM levels, while the NAD^+^ pool appears to be directly unaffected by NNMT. Through a complex metabolic perturbation, NNMT promotes cellular senescence and partial epithelial-to-mesenchymal transition, two key phenotypes that contribute to the progression of CKD.

We started by performing an unbiased analysis of publicly available human datasets, identifying NNMT as the gene most significantly negatively correlated with kidney function in CKD. Spatial transcriptomics using a novel segmentation strategy combining morphological markers with NNMT expression^63^, validated the association between tubular NNMT expression and senescence, showing that NNMT+ tubules interact more with their inflammatory microenvironment, to promote fibrosis. Additionally, we observed elevated renal NNMT expression in two preclinical models of early-stage CKD, with a strong correlation between NNMT and senescence at both the transcriptomic and protein levels. This increase occurred prior to fibrotic lesions in diabetic *db/db* mice and before renal functional decline in naturally aged mice, highlighting the early involvement of NNMT in these CKD models^42,43^. Further supporting this, we found elevated urinary levels of Me6PY, the end product of the NNMT reaction, in diabetic patients with microalbuminuria. Me6PY, a uremic toxin known to accumulate in the circulation during late-stage CKD, is excreted by the kidney^33,58^. In patients with early CKD, we did not observe impaired Me6PY elimination, but rather an overall increase in its levels compared to diabetic patients, suggesting a local increase in NNMT expression within the kidney, even before advanced disease stages. Similarly, preclinical models of early CKD exhibited decreased SAM levels and increased MeNAM levels locally in the kidney, further indicating altered NNMT activity in early stages of disease.

We also conducted *in vitro* mechanistic studies to decipher the functional role of tubular NNMT in the early stages of CKD. Overexpression of *Nnmt* in cultured tubular epithelial cells primed fibrosis and senescence without inducing a full detrimental phenotype. In contrast, stimulation with TGFβ, already shown to promote senescence and EMT^44,45^, functionally exacerbated NNMT-dependent fibrosis and senescence, in line with the worsening of renal histological lesions observed by Takahashi *et al.* in *Nnmt* transgenic mice subjected to ureteral obstruction^58^.

One likely explanation for the TGFb-mediated NNMT exacerbation, is the disruption of metabolic homeostasis by NNMT overexpression. NNMT catalyzes the conversion of NAM and SAM to MeNAM and SAH. NAM is a precursor of the essential cellular cofactor NAD^+^ and is incorporated into NAD^+^ by NAMPT, either after dietary intake or through intracellular recycling via the salvage pathway^64^. Kidney NAD^+^ levels are known to decline as CKD progresses^65^. Although NNMT overexpression is consistent with a model in which the excretion route diverts NAM from its NAD^+^ fate, our data and those of others indicate that NNMT is unlikely to directly regulate NAD^+^ levels in kidney disease. Our results indicate that NAD^+^ levels remain unchanged in *Nnmt*-OE cells. In addition, while TGFβ treatment in cells or hypoxia/reoxygenation damage in organoids reduced NAD^+^ levels, NNMTi did not restore NAD^+^ levels under these conditions. Similarly, the NAD^+^ pool was not altered when *Nnmt* was knocked down in mouse liver *in vivo* and in primary hepatocytes *in vitro*^66^. Furthermore, the affinity of NAMPT for its substrate is less than 1 µM^67^, which is higher than the affinity of NNMT for NAM, which is about 430 µM^68^. NAMPT is therefore considered to be saturated with NAM, and NNMT expression is unlikely to lower NAM to levels that would limit NAD^+^ synthesis^62^. In addition, we observed equilibrated NAM levels in the kidneys of *db/m* and *db/db* mice, as well as young and aged mice, despite increased NNMT expression. This is in line with studies showing no increase in NAM levels after knockdown of *Nnmt* in liver or adipocytes^66,69^. Thus, it is possible that NAM has a rapid intracellular or intratissular equilibration rate, explaining why NAM supplementation is not efficient in restoring tubular homeostasis in the context of NNMT overexpression. Indeed, our findings demonstrated no benefit of NAM supplementation on senescence and fibrosis in the aged preclinical model of CKD, and similarly, no improvement in kidney function was reported with short-term NAM treatment in *db/db* mice^70^. However, NAM supplementation has been shown to reduce kidney inflammation and fibrosis in more advanced preclinical disease models, such as unilateral ureteral obstruction^71^ and adenine-induced CKD^72^, although these studies did not report data on renal NNMT expression. However, we cannot exclude that NNMT expression may influence NAM/NAD^+^ metabolism in other tissues or metabolic conditionss^33,73^.

Conversely, our results show that SAM supplementation attenuates cellular senescence and EMT *in vitro*. In human CKD, urinary SAM levels have been shown to decrease in parallel with kidney function, making SAM a potential biomarker for monitoring kidney function decline in the early stages of the disease^74^. Consistently, we observed reduced SAM levels in the kidneys of *db/db* compared to *db/m* mice, as well as in aged mice compared to young. SAM is a universal methyl donor, required as a co-substrate for all methyltransferases, including DNA and histone methyltransferases. Because NAM methylation is irreversible, NNMT may act as a methyl donor sink, resulting in a reduction of epigenetic methylation marks and gene expression changes, a model originally implicated in cancer^75,76^. In the context of renal fibrosis, NNMT overexpression has been associated with DNA hypomethylation at the level of *Ctgf*, *p53* and *Sphk1* genes, resulting in a heightened expression^58,59,61^. While we did not detect changes in p53 expression in NNMT+ tubules from DKD kidney biopsies, our transcription factor enrichment analysis identified p53 as a likely contributor to the differentially expressed gene signature. Therefore, it is plausible that decreased SAM and methyl pool availability due to NNMT increased expression contribute to the early damage phenotype by altering regulation of p53.

While therapeutic strategies to decelerate CKD have emerged over the past decade, early detection and treatment of the root causes remain challenging^4^. Existing therapies for CKD primarily aim to control blood pressure, blood glucose, and inflammation, often involving treatments like renin-angiotensin system blockers or sodium-glucose cotransporter 2 inhibitors^77^. Although these interventions address key risk factors such as hypertension and hyperglycemia, they do not directly target the cellular and metabolic disturbances driving CKD ^5^. Increasing evidence suggests that targeting cellular senescence offers a promising early intervention for CKD, with senotherapeutic approaches successfully preventing fibrosis and kidney function decline^15,18^.

In this study, we demonstrate the benefits of NNMT inhibition in preventing the senescent and pro-fibrotic phenotype in damaged TECs and iPSC-derived kidney organoids by targeting a fundamental metabolic pathway involved in cellular energy homeostasis and epigenetic regulation. Importantly, the same selective and membrane-permeable NNMTi used in this study, 5-amino-1-methylquinolinium, has been successfully administered systemically in mice, improving muscle function in aged models^79^ and normalizing body composition and liver physiology in obese mice^80^. Preclinical pharmacokinetic and safety studies have validated NNMTi as a viable pharmacological approach with an acceptable off-target profile^21^, and it is currently under consideration for clinical trials targeting age-related muscle decline. In our study, we validated the use of a human-relevant kidney organoid model to successfully replicate tubular senescence and EMT, with NNMTi treatment normalizing these pathways in this clinically relevant system^78^. Collectively, our *in vitro* studies further support NNMTi as a potential senotherapeutic for age-related conditions.

In conclusion, our findings highlight the critical role of NNMT in promoting tubular senescence and fibrosis. NNMT inhibition thus emerges as a promising senotherapeutic strategy to preserve tubular homeostasis, mitigate early kidney function decline and slow CKD progression.

## METHODS

### Experimental models and subject details

#### Human DKD kidney specimen

Kidney tissue sections were derived from clinical biopsies collected at Amsterdam University Medical Center (UMC). Samples from ten male DKD patients were obtained from formalin-fixed paraffin-embedded (FFPE) tissue. The study was approved by the Amsterdam UMC Medical Research Ethics Committee, under the Biobank Assessment Committee number BTC 2015.081. Written informed consent was obtained from living donors. Information on sex, age, date of biopsy, eGFR and % IFTA is listed in **Figure S1F**.

#### Human DKD serum and urine samples

Serum and urine samples from diabetic and early DKD patients were obtained from the following clinical trials: Renal Oxygenation, oxygen Consumption and hemodynamic Kinetics in type 2 DIabetes: an Ertugliflozin Study (ROCKIES, NCT04027530), Renoprotective Effects of Dapagliflozin in Type 2 Diabetes (RED, NCT02682563), and Renal Actions of Combined Empagliflozin and LINagliptin in type 2 diabetES (RACELINES, NCT03433248). Early DKD was defined as 24-hour albuminuria > 30 mg. Samples collected at baseline were used. Baseline characteristics (sex, age, BMI, glycated hemoglobin A1c, fasting plasma glucose, duration of diabetes, albuminuria and eGFR) are provided in **Figure S1H**.

#### Animal studies

Adult male BKS.Cg-Dock7m +/+ Leprdb/J HE/HE (*db/m*) mice and adult male BKS.Cg-Dock7m +/+ Leprdb/J WT/HO (*db/db*) mice at 7 weeks of age at the time of receipt were purchased from Charles River (JAX Laboratory). Mice were housed individually with ad libitum access to water and food (normal chow diet, AIN-93M, Research Diets Inc.), until 16 weeks of age.

4 months old (young) and 21 months old (aged) wild-type (WT) C57BL/6JRj male mice were purchased from Janvier Laboratories. Mice were fed a Safe 150 diet (Safe Diets) and for the dietary supplementation, nicotinamide (NAM, Sigma-Aldrich #72340) was mixed with the diet at a dose of 200 mg/kg bw/day, 1.6g/kg food for 14 weeks.

Throughout both studies, mice were maintained at a standard temperature of 22 °C and in a humidity-controlled environment with a 12:12h light/dark cycle. Mice were housed with appropriate nesting material. At the end of the studies, animals were sacrificed via cardiac exsanguination followed by cervical dislocation, and urine and blood samples were collected. Immediately after, the kidneys were harvested, weighed and processed: one fixed in formalin for histology, and one snap-frozen in liquid nitrogen and kept in −80 °C for molecular biology readouts.

All procedures were performed in accordance with Swiss and EU ethical guidelines and approved by the Nestlé ethical committee, the French Ministry of Research (APAFIS#25310) and the Office Vétérinaire Cantonal Vaudois (VD3519x1).

#### Cell cultures

##### Renal tubular epithelial cells

TECs isolated from Immorto mice^81^ were maintained in DMEM/F12 medium (Thermo Fisher #21331020) with 10% fetal bovine serum (FBS, Thermo Fisher), 5 ug/ml insulin and transferrin, 5 ng/ml sodium selenite (Thermo Fisher #41400045), 40 pg/ml triiodo-thyrionine (Sigma-Aldrich #T5516), 36 ng/ml hydrocortisone (Sigma-Aldrich #H0135), 20 ng/ml epidermal growth factor (Sigma-Aldrich #E9644), 2mM L-glutamine (Thermo Fisher #25030081) and 1% penicillin/streptomycin (Thermo Fisher #15140122) and IFNγ (10 ng/ml, Prospec #CYT-358) at 33°C, 5% CO_2_. Before being seeded for experiments, cells were grown at restrictive conditions (37°C, 5% CO_2_, in the absence of IFNγ) for 7 days, to downregulate SV40 large T antigen activity. Cells were regularly tested using a Rapid Mycoplasma Detection Kit (MycoGenie, Assay Genie).

For cell transduction, lentiviral particles of pLenti-C-Myc-DDK-P2A-puro (PS100092V) or pLenti-mouseNNMT-MYC-DDK-P2A-puro (MR203499L3V) were purchased from Origene. TECs were incubated for 18 hours with lentiviral particles at 33°C, 5% CO_2_. Cells were subsequently selected for 7 days with 5 µg/ml puromycin.

For experimental readouts, cells were shifted to DMEM/F12 medium with 10% FBS, glutamine and antibiotics, plus the following compounds as appropriate: Transforming Growth Factor-Beta 1 (TGFβ, Prospec #CYT-716), 5-amino-1-methylquinolinium iodide (NNMTi, Sigma-Aldrich #SML2832), nicotinamide (NAM, Sigma-Aldrich #72340), and S-(5′-Adenosyl)-L-methionine iodide (SAM, Sigma-Aldrich #A4377). TECs were stimulated 48h with 10 ng/ml TGFβ, 50 µM NNMTi, 1 mM NAM and 0.5 mM SAM. Optimal concentrations were chosen based on MTT cell viability assay (Roche #11465007001).

##### iPSC-derived kidney organoids

Human induced pluripotent stem cells (iPSCs) were kindly provided by J. Jansen (Radboud University Medical Center, Nijmegen, the Netherlands). Briefly, human adult skin fibroblasts derived from a healthy volunteer were reprogrammed into iPSCs using the Yamanaka factors by the Stem Cells Technology Center at Radboud University Medical Center (SCTC, Radboud UMC, Nijmegen, The Netherlands).

iPSC colonies were cultured on Geltrex (Thermo Fisher #A1413301) coated 6-well plates (Greiner Bio-one) in StemFlex medium (Thermo Fisher #A3349401), and split using EDTA (Thermo Fisher #15575020). To make kidney organoids, iPSCs were seeded at 180’000 cells per well in a coated 6-well plate. The differentiation protocol was based on Takasato *et al.* and Jansen *et al.*^51,82^. On day 0 (d0), differentiation was initiated using CHIR-99021 (6 μM, R&D systems #4423) in STEMdiff APEL2 medium (Stemcell Technologies #05270). CHIR treatment was maintained for 3 (differentiation towards ureteric bud) and 5 days (differentiation towards metanephric mesenchyme), after which the medium was supplemented with fibroblast growth factor 9 (FGF9, 200 ng/ml, Peprotech #100-23) and heparin (1 μg/ml, Sigma-Aldrich #H4784) up to d7. On d7, cell layers were trypsinized with 0,05% trypsin-EDTA (Thermo Fisher #25300) and 3 days CHIR-differentiated cells were mixed with 5 days CHIR-differentiated cells to a ratio of 1:2. To generate cell aggregates, cells were aliquoted in a 1.5-ml tube at a density of 300’000 cells per tube. The cells were then subjected to 3 cycles of centrifugation at 300 rcf for 3 minutes, with a change in position by 180° between cycles. Cell aggregates were plated on Costar Transwell filters (Corning #3450) and cultured at an air-medium interface, thus creating a 3D organoid model. A one-hour CHIR pulse (5 μM) in APEL2 medium was used to stimulate self-organizing nephrogenesis and medium was replaced for APEL2 supplemented with FGF9 and heparin for an additional 5 days. As of d7+5, organoids were cultured using Essential 6 medium (E6, Thermo Fisher #A1516401) supplemented with human epidermal growth factor (hEGF, 10 ng/ml, Sigma-Aldrich #E9644), bone morphogenetic protein-7 (BMP7, 50 ng/ml, Peprotech #120-03P) and stromal derived factor 1 beta (SDF1β, 10 ng/ml, R&D systems #351-FS) until further experimental procedures at d7+17.

For hypoxia/reoxygenation experiments, d7+17 organoids were incubated in the HypOxystation H35 (Don witley scientific) in 1% Oxygen, 5% CO_2_ at 37°C. Normoxic controls were kept in culture in E6 medium without any growth factor. After 2 days of hypoxia, both hypoxic and normoxic organoids received fresh medium, supplemented with 50 µM NNMTi if applicable, and were cultured for an additional 24 h under standard conditions (re-oxygenation).

### Method details

#### Bioinformatics analyses of human publicly available datasets

Datasets were selected by screening the Nephroseq (v5) database considering microarray transcriptomics datasets with available eGFR values. Datasets were filtered on the existence of tubulointerstitium samples and included if they contained >2 samples with a Diabetic Kidney Disease phenotype. Three datasets were considered: two from GSE47184 (Affymetrix Human Genome HG-U133A Custom CDF: dataset 1, and Affymetrix Human Genome U133 Plus 2.0 Array: dataset 2)^22^, and Affymetrix Human Genome U133A 2.0 Array (dataset 3) data from GSE30529^23^.

Before merging the datasets, the raw data was pre-processed. The CEL files were downloaded from the OMIM repository and individually checked for outliers through calculating Relative Log Expression (RLE) and Normalized Unscaled Standard Errors (NUSE). Samples that showed a NUSE over 5%, were considered outliers and excluded from the analysis. This led to the exclusion of 1 sample from both the datasets 2 and 3. The gene chips were background corrected, normalized and probes were summarized by fitting a GeneChip Robust Multiarray Averaging method (GC-RMA) using the R/Bioconductor package gcrma version 2.70.0. Further outliers were determined in each dataset as sample points that were outside of the 95% confidence interval ellipse of a Principal Component Analysis (PCA), determined by calculating the Mahalanobis distance of each sample from the center of the PCA plot. A sample was considered an outlier if the Mahalanobis distance was larger than the chi-square critical value at the 95% confidence level. This led to the further exclusion of 2 samples from each of the datasets 1, 2 and 3. To correct for batch effects, we used the ComBat algorithm^83^, which was implemented with the sva version 3.46.0 package. To ensure comparison between the different datasets, we identified common Entrez gene identifiers across the different platforms and merged the datasets using only these common identifiers. The resulting merged dataset had 104 samples and 6604 features.

Linear models were fitted to the data to assess the correlation between the gene expression levels and the eGFR for each gene. The significance of the correlation was assessed by Spearman’s correlation test. To assess the impact of CKD pathology on the correlation between *NNMT* gene expression and eGFR, linear regression models were employed with eGFR as the primary predictor and donor pathology as a covariate. Enrichment analyses were performed using fgsea implemented in clusterProfiler 4.6.2^84^ and the Molecular Signatures Database (MSigDB) v2023.2Hs collection H (hallmark gene sets). Genes were ranked according to their Spearman correlation coefficient with eGFR or *NNMT* expression. For the custom GSEA, we tested three validated senescence gene sets, SenMayo^25^, CSGene^26^ and Fridman_up^27^, and 2 gene sets related to renal tubulointerstitial fibrotic damage, the renal fibrosis gene set from DisGeNET^85^ and the epithelial-to-mesenchymal transition (EMT) hallmark from MSigDB v2023.2.Hs. P-values were corrected for multiple testing using the Benjamini-Hochberg correction.

#### Histology and immunohistochemistry of human and mice kidney tissue

Human kidney tissue sections (4 µm) were deparaffinized in xylene and rehydrated in ethanol. For interstitial fibrosis and tubular atrophy (IFTA) scoring, biopsy sections were stained with Jones’ Methenamine Silver Stain using a standard procedure on Ventana Benchmark Ultra (Roche Diagnostics). The percentage of IFTA was scored by a pathologist (J.J.T.H.R.) in a blinded fashion. For p21 and NNMT staining, slides were boiled for 10 minutes in Tris/EDTA buffer at pH 9 for antigen retrieval and incubated overnight with rabbit anti-human p21 (1:500, clone 12D1, CST #2947) and rabbit anti-human NNMT (1:3000, Proteintech #15123-1-AP) antibodies. Sections were then incubated with BrightVision goat anti-rabbit/polyHRP (Immunologic) secondary antibody for 30 min and subsequently stained with 3,3-diaminobenzidine (DAB). Nuclei were counterstained with hematoxylin. Slides were mounted using VectaMount mounting medium (Vectorlabs) and scanned with Philips IntelliSite Ultra-Fast Scanner. Images were analyzed using QuPath software^86^: p21 positive nuclei in the tubulointerstitial area were quantified using the *positive cell detection* command and NNMT positive areas were detected with a *pixel classifier*. For image overlays, sections were aligned using Philips Registration multiple batches program and images were stacked using FIJI software version 1.54f^87^.

Mouse tissue was fixed in a buffered 10% formalin solution (Sigma-Aldrich # HT501128) for 24 hours, dehydrated and embedded in paraffin. Three-micrometer sections were mounted on adhesive slides, and subsequently deparaffinized and rehydrated through xylene and ethanol washes. Slides were stained for Picro-Sirius-Red (PSR) and Periodic acid–Schiff–diastase (PAS-D) according to standard protocols and scanned using a Zeiss AxioScan.Z1 HF slide scanner (for the *db/db* mice study) or an Olympus VS120 slide scanner (for the aging mice study). Quantification of the relative proportion of fibrosis (PSR-positive area) was performed using Whole Slide Imaging (WSI) with the Visiopharm® software (for the *db/db* mice study) or the QuPath software (for the aging mice study). The IFTA score of the aging mice study kidneys was assessed by a pathologist (J.J.T.H.R.) in a blinded fashion on PAS-D-stained sections.

#### Spatial transcriptomics of human kidney biopsies

Spatial transcriptomics was performed on six DKD kidney biopsies using the NanoString GeoMx® Digital Spatial Profiler (DSP). Slides were prepared according to GeoMx DSP slide preparation user manual (MAN-10087-04). Briefly, FFPE tissue sections of 5-µm were mounted on positively charged histology slides and incubated at 60 °C for 1 h. Slides were deparaffinized in xylene (3 times 5 min) then rehydrated in ethanol gradient (2 baths of 5 min in 100% EtOH, followed by 5 min in 95% EtOH), and wash with PBS. Antigen retrieval was performed in Tris-EDTA pH 9.0 buffer at 100°C for 15 min at low pressure. Slides were then washed in PBS 2x, and incubated in 1 µg/ml proteinase K (Thermo Fisher #AM2548) for 15 min at 37 °C and washed again with PBS. Tissues were post-fixed in 10% neutral-buffered formalin for 5 min, washed twice for 5 min in NBF stop buffer (0.1 M Tris Base, 0.1 M Glycine), and finally once in PBS.

The RNA probe mix (GeoMx Human Whole Transcriptome Atlas, NanoString #121401102), a pool of in situ hybridization probes with UV photocleavable oligonucleotide barcodes, was placed on each section and covered with HybriSlip Hybridization Covers. Slides were then incubated for hybridization overnight at 37 °C in a Hyb EZ II hybridization oven (Advanced Cell Diagnostics). The day after, HybriSlip covers were gently removed and 25-min stringent washes were performed twice in 50% formamide and 2× Saline Sodium Citrate (SSC) at 37 °C. Tissues were washed for 5 min in 2× SSC, then blocked in Buffer W (provided in the GeoMx RNA Slide Prep FFPE-PCLN kit, NanoString #121300313) for 30 min at room temperature in a humidity chamber. Next, tissues were stained for tubular markers, i.e. AF647-panCK (1:100, Novus Biologicals #NBP2-33200) and AF488-megalin (1:100, Santa Cruz Biotechnology, clone H10 #sc-515772), the nuclear marker SYTO 13 (500nM, Thermo Fisher #S7575), and AF594-NNMT (1:50, Novus Biologicals #NBP2-72984) in Buffer W for 1 h at room temperature. Slides were washed twice in fresh 2× SSC and then loaded on the GeoMx DSP.

For GeoMx DSP sample collections, entire slides were imaged at ×20 magnification and the morphologic markers were used to select NNMT-positive tubules (NNMT+) and NNMT-negative tubules (NNMT-) regions of interest (ROIs) using organic shapes. Automatic segmentation of ROIs to define areas of illumination (AOIs) was based on tubular markers and NNMT staining. This allowed to separate tubules and surrounding cells (microenvironment). AOIs were exposed to 385 nm light (UV), releasing the indexing oligonucleotides that were collected with a microcapillary and deposited into a 96-well plate for subsequent processing. The indexing oligonucleotides were dried overnight and resuspended in 10 μl of DEPC-treated water.

Sequencing libraries were generated by PCR from the photo-released indexing oligos and AOI-specific Illumina adapter sequences, and unique i5 and i7 sample indexes were added. PCR reactions were pooled and purified twice using AMPure XP beads (Beckman Coulter #A63881). Pooled libraries were paired-sequenced at 2 × 27 base pairs and with a unique dual indexes workflow on an Illumina NovaSeq instrument. NovaSeq-derived FASTQ files for each sample were compiled for each compartment using the bcl2fastq program of Illumina, and then demultiplexed and converted to digital count conversion (DCC) files using the GeoMx DnD pipeline (v.2.0.0.16) of NanoString. Quality control was performed using the R/Bioconductor package GeomxTools v3.4.0. Each AOI was controlled for number of raw reads and sequencing quality and probes were checked for outlier status. AOI and/or genes with abnormally low signal were filtered out based on the distribution of negative probes and the percentage of genes detected per segment and of segments detecting a gene, both set at a minimum of 5%. Then, after ensuring a good separation between the upper quartile (Q3) of the counts in each AOI with the geometric mean of the negative control probes, AOI were normalized by the geometric mean of Q3 counts across all AOIs. The resulting dataset contains 12,062 genes.

Differential expression of gene targets was determined on Q3 normalized data via a mixed effects linear regression model using the R package lme4 v1.1-35.1. The regression model implemented was as follows: log2(gene) ∼ region + (1 | slide). Gene set enrichment analyses were performed using fgsea implemented in clusterProfiler 4.6.2^84^ and the MSigDB v2023.2Hs collections H (hallmark gene sets) and Reactome. Senescence gene sets enrichment analysis was performed using the same gene sets described above. To perform cell-type deconvolution, we used NanoString’s algorithm SpatialDecon 1.8.0^88^. Cell-type proportions were determined by comparing via log-normal regression our spatial transcriptomics data to a cell profile matrix derived from single-cell RNA-seq data of the human kidney^40^. For the visual representation, specific gene expression or calculated scores were plotted on the image data using the R/Bioconductor packages singscore 1.18.0 and SpatialOmicsOverlay v1.2.1. To predict the transcription regulators of differentially expressed genes (defined as adjusted p-value <0.05), we used the online version of Lisa^36^, which creates a chromatin landscape model based on Cistrome Data Browser^37^, and probes this model with publicly available ChIP-seq data. Lastly, cell-cell communication inference analysis was performed with CellChat^41^ and the CellChatDB.human library of receptor-ligand interactions. For this analysis, the transcript counts of one receptor gene in one AOI are combined with the corresponding ligand gene counts in a different AOI to create an interaction strength score.

#### Clinical chemistry analyses of mice

For the *db/db* mice study, GFR was measured with a transdermal monitor device. Briefly, at basal, mid and end study, animals were equipped with a miniaturized fluorescence detector directly attached to the skin on the back which measured the excretion kinetics of the exogenous GFR tracer, fluorescein-isothiocyanate (FITC) conjugated sinistrin (an inulin analog). Mice received an intravenous injection of 50mg/kg FITC-Sinistrin and the GFR was recorded for 1.5 hour on conscious, freely moving animals. Urinary albumin and creatinine levels were quantified with the Albuwell M (Albumin, Microalbuminuria), Mouse / Indirect Competitive Ab Capture ELISA Kit (Exocell #1011) and the QuantiChrom™ Creatinine Assay Kit (BioAssay Systems, # DICT-500), respectively, on end of study urine.

For the aging mice study, analyses were done with the Architect Ci4100 analyzer (Abbot Laboratories) with the clinical chemistry module C4000. The following kits from Abbot Laboratories were used: 2K98-24 for urinary albumin and 4S95-20 for plasma and urinary creatinine.

#### Bulk transcriptomics of cells and mice kidney tissue

The transcriptome of cellular and mice experimental models was measured with QuantSeq 3′ RNA sequencing. Total RNA was extracted from cells seeded in 6-well plates or from 10-20 mg of kidney tissue using Agencourt RNAdvance Tissue kit (Beckman Coulter Life sciences #A32646). RNA quantification was performed with the Quant-iT RiboGreen RNA Assay Kit (Thermo Fisher #R11490) and 100 ng was used to generate libraries using the QuantSeq 3’ mRNA-Seq V2 Library Prep Kit FWD with UDI (Lexogen #193.384). After 19 cycles of amplification, the libraries were quantified using Quant-iT PicoGreen dsDNA Assay Kit (Thermo Fisher #P7589). Size pattern was controlled with the high sensitivity NGS Fragment Analysis kit on a Fragment Analyzer (Agilent #DNF-474). Libraries were then pooled at an equimolar ratio and clustered at a concentration of 650 pM on a sequencing flow cell (Illumina). Single-read sequencing was performed for 65 cycles on a Nextel 2000 (Illumina) using the P3 50 kit (Illumina #20046810). Demultiplexing was performed on sequencer using Dragen version 4.2.7. Sequencing reads were trimmed with BBDuk (BBTools version 36.84, Bushnell B., sourceforge.net/projects/bbmap/). Mapping to the mouse genome (built GRCh38.101) was performed with RNA STAR^89^, version 2.7.3a, using default parameters. Gene count was performed with HTSeq^90^, version 2.16.2.

Using the R/Bioconductor packages limma 3.54.2^91^ and edgeR 3.40.2^92^, raw counts were converted to counts per million (CPM) and lowly expressed genes were filtered out by selecting only genes with at least 15 reads in a minimum number of samples, corresponding to the minimum group sample size. The filtering step was performed on the CPM values to take into account the library size. The resulting dataset was then normalized by the method of trimmed mean of M-values (TMM). For the statistical comparison of expression changes, we fit a linear model and constructed a contrast matrix to assess differential expression between groups. GSEA was then performed using CAMERA^93^ and the MSigDB v2023.2.Mm collection MH (orthology-mapped hallmark gene sets). For selected pathway enrichment, we used fry, which performs a fast approximation of the roast algorithm (rotation gene set testing), and the R/Bioconductor package clusterProfiler 4.6.2 for data visualization. We specifically tested for the mouse versions of SenMayo^25^ and CSGene^26^, the mouse orthologs of DisGeNET renal fibrosis^85^ and the epithelial-to-mesenchymal transition hallmark from MSigDB v2023.2.Mm. To perform GSEA of *Nnmt* correlated genes, we ranked the genes by Pearson correlation coefficient to *Nnmt* expression and used clusterProfiler fgsea function.

#### Metabolomics of human serum and urine

Serum and urine samples (60 µL) were extracted with 800 µL of 80% (v/v) cold methanol containing internal standards (5 µM NAM-d4 and trigonelline-d3). Samples were vortexed and incubated at 4 °C for 30 min, under agitation, before being centrifuged for 15 min, and the supernatant was collected. The supernatant was evaporated using a vacuum centrifuge. The dried extracts were stored at −80 °C until analysis. Prior to analysis, the extracts were reconstituted in 60 µL (urine) or 120 µL (serum) of 60% (v/v) acetonitrile/water and transferred to glass vials. The analysis of NAD^+^ and related metabolites in serum and urine was performed using a method adapted from Giner *et al.*^94^, with a few modifications. The used ultra-high-performance liquid chromatography (UHPLC) system consisted of a binary pump, a cooled autosampler, and a column oven (1290 Infinity II LC System, Agilent), coupled a time-of-flight mass spectrometer (ZenoTOF 7600 System, Sciex) equipped with a Turbo V ion source, operating in positive electrospray ionization (ESI) mode, by acquiring multiple reaction monitoring (MRM) for each metabolite. In comparison to the cited method, NMNH and NR-d4 were not monitored. Acquisition and data analysis was controlled by SCIEX OS-Q version 3.1.6.44 (Sciex).

#### Metabolomics of cells and mice kidney tissue

Liquid-liquid extraction was performed as described elsewhere^94^. Cells were scraped on dry ice from 6-well plates after adding 800 µl −20°C methanol:water (5:3 (v/v)) including isotopically labelled internal standards. After transferring to a tube, 500 µl −20°C chloroform were added and the samples were homogenized in a shaker (Thermomixer C, Eppendorf) at 4°C for 10 min. Tissue samples of ∼10 mg were extracted with 1.3 mL −20°C methanol:water:chloroform (5:3:5 (v/v)) containing isotopically labelled internal standards. The samples were transferred to pre-cooled racks (-80°C) of a tissue mixer and homogenized with the aid of metal beads for 3 rounds, 3 min, 28 hz. After homogenization, the cell and tissue samples were centrifuged at 21130 rcf at 4°C for 10 min and the resulting polar and apolar phases were transferred into a new tube. The remaining protein disc was dried, solubilized in 0.2 M KOH for 20 min at 90°C, and quantified with a BCA assay (Thermo Fisher) for later data normalization. 300 µl of each extract was pooled to generate quality control samples. The extracts and the quality control samples containing different amounts of the pooled extract (100-1500 µl) were dried overnight in a vacuum centrifuge (Labconco) at 4°C and 4 mbar. Prior to measurement the dried samples were resuspended in 35 µl 70% (v/v) acetonitrile:water.

Metabolite separation and detection was achieved with liquid chromatography (LC) and high-resolution mass spectrometry (HRMS), respectively, consisting of Vanquish LC system (Thermo Fisher) connected to a Lumos Orbitrap system (Thermo Fisher). For chromatographic separation a ZIC-pHILIC column (100 x 2.1 mm, 5 µM, Merck) with a guard-column (20 x 2.1 mm, 5 µM; Merck) was used. The two mobile phases were A) 10 mM ammonium acetate containing 0.4% (v/v) ammonium hydroxide (pH∼9.3) and B) acetonitrile. The metabolites eluted over a linear gradient from 90-25% B in 16 minutes at 35°C. The mass spectrometer was operated in full scan (m/z: 72-950) at a resolution of 50’000 at an m/z of 200 with alternating positive and negative mode acquisitions. The positive and negative mode voltages were 3500 V and 3000 V, respectively, the ion transfer tube was 310 °C, and the vaporizer temperature was 280° C. The sheath gas was set to 30 and the auxiliary gas to 15. The metabolites were integrated based on their accurate mass and retention time in a semi-targeted fashion with Xcalibur (Thermo Fisher). The metabolites were normalized by the best fitting internal standard, which was determined with the QC samples according to linearization, and protein content.

#### Western blot analyses of cells and mice kidney tissue

Cells were scrapped from 6-well plates in RIPA buffer (Sigma-Aldrich #R0278) supplemented with “cOmplete, Mini, EDTA-free Protease Inhibitor Cocktail” (Roche, Sigma-Aldrich #11836170001) and “PhosSTOP” (Roche, Sigma-Aldrich #4906837001). 30-40 mg of kidney tissue was lysed in RIPA buffer with protease and phosphatase inhibitors using a Polytron homogenizer (Kinematica). After a centrifugation step, the protein content of the clear supernatant was quantified with a BCA assay kit (Thermo Fisher #23225). Protein extracts were denatured in LDS Sample Buffer (Thermo Fisher #NP0007) and Sample Reducing Agent (ThermoFisher #NP0009) at 70°C for 10 minutes. 10-20 µg of protein was separated by SDS-PAGE on 4–12% gradient gels (Thermo Fisher #NP0336BOX and #WG1403BOX) with MES SDS Running Buffer (Thermo Fisher #NP0002) and transferred onto PVDF membranes using an iBlot 2 Gel Transfer Device and appropriate transfer stacks (Thermo Fisher #IB24001). Membranes were blocked in 5% non-fat dry milk in Tris-buffered saline-Tween 20 (TBST, Thermo Fisher #28360), and incubated overnight at 4°C with primary antibodies anti-p21 (1:500, Abcam #107099, kidney tissue), anti-p21 (1:500, Abcam #188224, cells), anti-NNMT (1:500, Abcam #119758, kidney tissue), anti-NNMT (1:500, Proteintech #15123-1-AP, cells), anti-Twist (1:1000, Cell Signaling #90445, cells), anti-Snail (1:1000, Cell Signaling #3879S, cells), and anti-Vinculin (1:1000, Cell Signaling #13901, kidney tissue and cells). Membranes were then washed and incubated for 1h with horseradish peroxidase–conjugated donkey anti-rabbit, anti-mouse and anti-rat (1:3000, Jackson ImmunoResearch, #711-035-152, #715-035-150, #712-035-150). Proteins were visualized with chemiluminescent western blotting substrate (Thermo Fisher, #32209 and #34096) using e-Blot Touch Imager. Densitometry analysis was performed using FIJI version 1.54f^87^. Protein levels in each lane were normalized to vinculin levels on the same gel, used as a loading control.

#### Flow cytometry analyses of cells

At the end of the experimental treatment, cells seeded in 12-well plates were incubated 1h at 37°C with 10 µM EdU (part of Click-iT Plus EdU Alexa Fluor 647 Flow Cytometry Assay Kit, Thermo Fisher #C10635). Cells were trypsinized and fixed in 4% paraformaldehyde. Between each post-fixation step, samples were washed with 1% BSA in phosphate-buffered saline (PBS, Thermo Fisher #10010056). Cells were stained 1h in an incubator at 37°C without CO_2_ with the CellEvent Senescence Green Flow Cytometry Assay Kit (1:1000, Thermo Fisher #C10840), and subsequently permeabilized in Triton-X 0.1% in PBS for 15 minutes. The Click-iT reaction was then run for 30 minutes. Samples were stored at 4°C and on the day of data acquisition, cells were stained with DAPI at 1 µg/mL and incubated for 30 minutes. DAPI-stained samples were directly analyzed on a 5-laser (355, 405, 488, 561 and 640 nm) Becton Dickinson LSRFortessa flow cytometer using the following parameters for excitation and emission: DAPI (355 nm laser, 450/50 nm detector), CellEvent Senescence (488 nm laser, 525/50 nm detector), EdU (640 nm laser, 660/20 nm detector). All fluorescence-bearing reagents were titrated, and their relative incubation time was optimized. Spectral overlap between fluorophores was corrected by compensation using single-stained controls. Data analysis was performed with FCS Express version 7.

#### RT-qPCR analyses of cells and organoids

RNA from cells was isolated using TRIzol reagent (Thermo Fisher #15596026), chloroform and isopropanol precipitation, and RNA from organoids was isolated using the PureLink RNA Mini Kit (Thermo Fisher #12183018A) according to the manufacturer’s protocol. cDNA was synthesized using M-MLV reverse transcriptase (Fisher Emergo #10247352) and oligo-dT primers. Transcript analysis was performed by real-time quantitative PCR on the Roche Light Cycler 480 using SYBR green master mix (Bioline #CSA-01190). Relative gene expression was analyzed using LinRegPCR version 2021.2^95^. Gene expression was normalized to human Tata Box binding Protein (TBP) housekeeping gene. The following primer sequences were used: *Nnmt*, forward: 5’-TGTGATCTTGAAGGCAACAGA-3’, reverse: 5’-CTTGATTGCACGCCTCAAC-3’; *NNMT*, forward: 5’-TGTGTGTGATCTTGAAGGGAAC-3’, reverse: 5’-CTTGACCGCCTGTCTCAAC-3’; *CDKN1A* (p21), forward: 5’-GACTCTCAGGGTCGAAAACG-3’, reverse: 5’-GGATTAGGGCTTCCTCTTGG-3’; *MCP1*, forward: 5’-CTGCTCATAGCAGCCACCTT-3’, reverse: 5’-GCACTGAGATCTTCCTATTGGTG-3’; *IL6*, forward: 5’-CTGCAGCCACTGGTTCTGT-3’, reverse: 5’-GATGAGTACAAAAGTCCTGAT-3’; *CTGF*, forward: 5’-AGCTGACCTGGAAGAGAACATT-3’, reverse: 5’-GCTCGGTATGTCTTCATGCTG-3’; *ACTA2* (αSMA), forward: 5’-ACTGCCTTGGTGTGTGACAA-3’, reverse: 5’-CACCATCACCCCCTGATGTC-3’; *COL1A1*, forward: 5’-AGGTGAAGCAGGCAAACCT-3’, reverse: 5’-CTCGCCAGGGAAACCTCT-3’; *FN1*, forward: 5’-GCGAGAGTGCCCCTACTACA-3’, reverse: 5’-GTTGGTGAATCGCAGGTCA-3’; *TGFΒ1*, forward: 5’-CCTGCTGTACTGCGTGTCCA-3’, reverse: 5’-CCAACTATTGCTTCAGCTCCA-3’; *SOX9*, forward: 5’-AGCGAACGCACATCAAGAC-3’, reverse: 5’-CTGTAGGCGATCTGTTGGGG-3’.

#### Whole mount organoid staining and imaging

Organoids were harvested from the transwell with a scalpel blade and transferred to 24-wells plate containing PBS. Organoids were washed twice with DPBS (Gibco) and fixed with 2 % PFA for 20 minutes at room temperature. Subsequently, organoids were washed twice with DPBS and stored in DPBS+0.2% PFA at 4° C until needed for staining. On the day of staining organoids were washed 3x for 10 minutes at room temperature with DPBS while gently shaking. Organoids were blocked with DPBS + 10% donkey serum (Jackson ImmunoResearch) + 0,06% Triton X-100 for 2 hours at room temperature while gently shaking. Organoids were incubated with primary antibodies (LTL-biotin, Vector laboratories #B-1325; NNMT, Abcam #119758; p21, Cell Signaling #2947) diluted 1:300 in DPBS + 10% donkey serum + 0,06% Triton X-100 overnight at 4° C. Next, organoids were washed 3x with DPBS + 0.3 % Triton X-100 and incubated with secondary antibodies (Streptavidin AF405, Invitrogen #S32351; Donkey a-mouse IgG H+L AF488, Invitrogen #A-21202; Donkey a-Rabbit IgG H+L AF647, Invitrogen #A-31573) diluted 1:400 in DPBS at room temperature for 2 hours. Subsequently, organoids where washed 3x with DPBS and mounted using Aqua-Poly/Mount (Polysciences). Organoids were imaged using a Stellaris 5 (Leica). Images were processed using LAS X.

#### NAD^+^ quantification in organoids

NAD^+^/NADH total pool was quantified by a colorimetric assay using the NAD/NADH Quantification Kit (Sigma-Aldrich #MAK037), according to the manufacturer’s protocol. NAD concentrations were normalized to the protein content of organoids using a BCA assay kit (Thermo Fisher #23225).

#### Mitochondrial respiration assay in cells

Cellular oxygen consumption rate (OCR) was determined using a Seahorse XFe96 analyzer (Agilent Technologies). Briefly, cells were seeded in XFe96 cell culture microplates (Agilent Technologies #103794-100) coated with 0.1% gelatin (Abm #TM063). At the end of the experimental treatment, medium was exchanged for Seahorse XF DMEM medium pH 7.4 (Agilent Technologies #103575-100) supplemented with 15 mM glucose (Thermo Fisher #A2494001), 1 mM sodium pyruvate (Thermo Fisher #11360070) and 2 mM glutamine (Thermo Fisher #25030081). Cells were allowed to equilibrate in a CO_2_-free incubator at 37°C for 30 minutes before being placed in the analyzer. During the metabolic assay, OCR rates were measured over time at basal levels and after the sequential addition of respiratory chain inhibitors and uncouplers at the following concentrations: oligomycin (2.5 μg/ml), FCCP (2.5 μM) and antimycin A (1 μg/ml) along with rotenone (1 μm). Cells were lysed with 1% SDS, 0.1 N NaOH, and protein content was measured using a BCA assay kit (Thermo Fisher #23225) for normalization of respiration measurements in each well.

#### Statistics

All statistical analysis and data visualization was carried out using GraphPad Prism 10.2.2 or R version 4.2.2 along with the tidyverse package^96^. The packages and statistical methods used for omic analyses are described in the respective methods sections. For the other experimental readouts, all data were tested for normality using the Shapiro-Wilk test. When comparing two conditions, Student’s two-tailed unpaired t-test was used. In case of unequal variance, Welch’s correction was applied. For comparisons of more than two groups, data were analyzed using one-way analysis of variance (ANOVA) followed by Šídák’s post hoc test for multiple comparisons, except in the case of ordinal data, for which the nonparametric Kruskal-Wallis test with Dunn’s post-hoc test was used. P values of <0.05 (*), <0.01 (**), <0.001 (***) and <0.0001 (****) were considered statistically significant. n.s. indicates results that were not statistically significant. Data are expressed as mean±standard error of the mean (s.e.m.). The statistical methods used for each analysis are mentioned in the figure legends.

## Data availability

Bulk transcriptomics data from mouse kidneys and cells will be available upon publication.

## ACKNOWLEDGEMENTS

We thank Carles Canto (EPFL), the NIHS community, and the members of the nephropathology research group at the Amsterdam UMC for critical discussion of the results. We also thank Alix Zollinger (Nestlé Research) for discussions on statistics and data visualization, Benjamin Brinon (Nestlé Research) for support with the metabolomics extraction, Jitske Jansen and Martijn van den Broek (Radboud UMC) for sharing their valuable insights on kidney organoids culture, the EPFL Histology Core Facility (HCF) and the Bioimaging and Optics Platform (BIOP) for histology and image analysis technical support, the CHUV Immune Landscape Laboratory Platform (ILL) for the spatial transcriptomics experiments, and Biomeostasis for the *db/db* preclinical study. The project was funded by Nestlé Research and direct funding to the senior authors. V.S.’s research is supported by the NUHS Internal Grant Funding under NUS Start-up grant NUHSRO/2022/047/Startup/11, an MOE Tier 2 grant T2EP30124-0014 and a Vidi grant from the Netherlands Organization for Scientific Research (NWO; 09150172110059). AT’s research is supported by the NWO–FAPESP joint grant on healthy ageing, executed by the Dutch research council (Zonmw 457002002).

## AUTHOR CONTRIBUTIONS

L.C., M.J.S., J.N.F., V.S., and A.T. designed the experimental strategy, interpreted the results and led the project. L.C., V.S., and A.T. wrote the manuscript. L.C., H.L., L.B., and S.L. performed *in vitro* experiments and analyzed data. L.C. and C.W. carried out the bioinformatics analyses. J.T., S.M., and J.A.H. performed (spatial) transcriptomics experiments and supported the bioinformatics. L.C. and L.T. carried out and analyzed flow cytometry experiments. L.C. and N.C. performed and analyzed human histology. L.C., S.C., V.F., and S.M. conducted and analyzed metabolomics. L.C., S.K., G.L., and J.L.S.-G. carried out the murine *in vivo* studies. E.J.M.v.B., M.J.B.v.B., A.C.H., D.v.R., and J.J.T.H.R. provided access to clinical samples. All authors read and edited the manuscript.

## SUPPLEMENTARY FIGURES

**SUPPLEMENTARY FIGURE 1.**
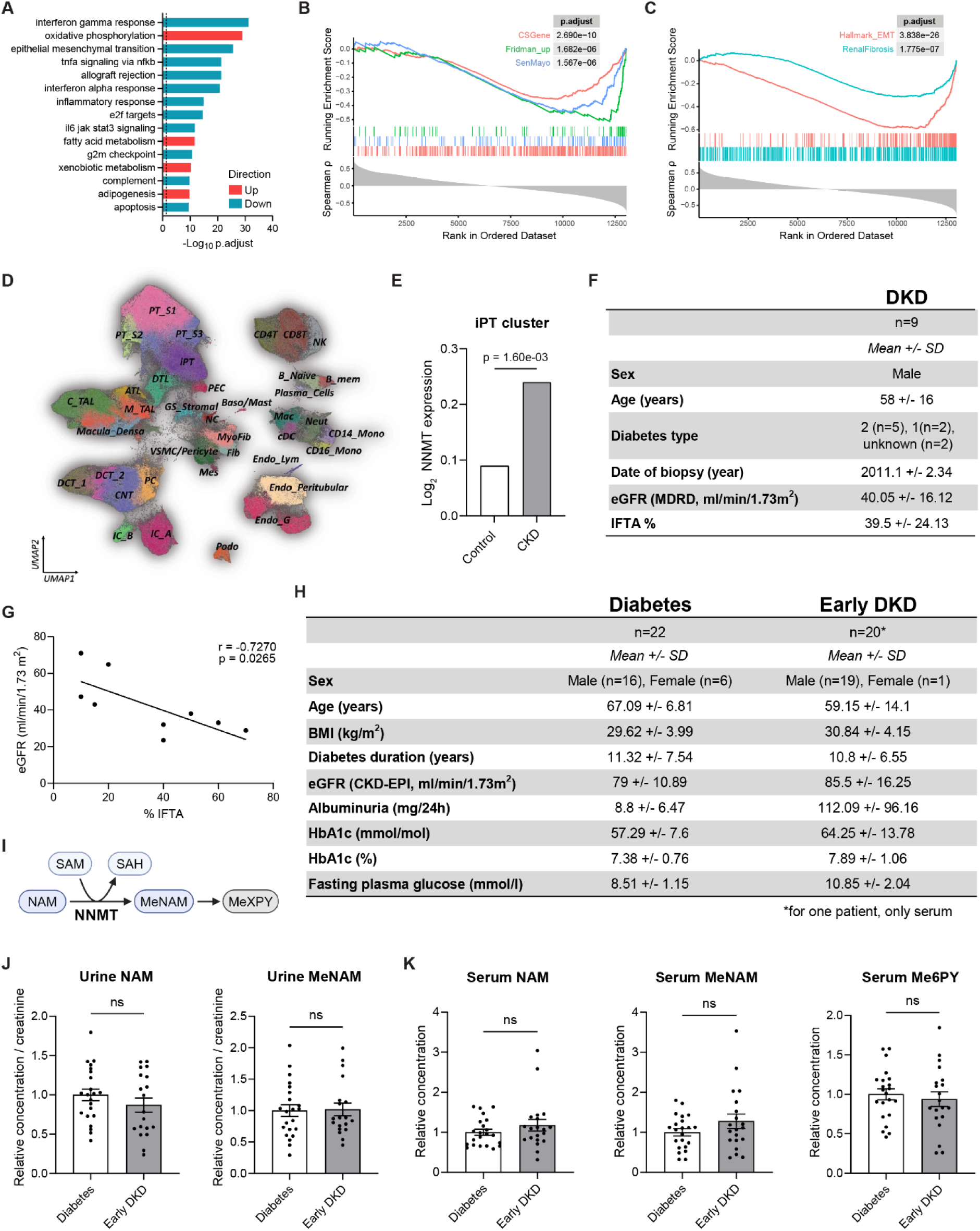
Tubular senescence and fibrosis are hallmarks of kidney dysfunction and associate with tubular NNMT expression. **(A)** Transcriptome associated with eGFR in CKD patients. Top 15 gene sets with the lowest p.adjust using GSEA of the hallmark curated gene set collection from MSigDB. n=104 from the combined dataset. **(B,C)** GSEA plots, ranking genes by Spearman correlation coefficient to eGFR and showing enrichment in (B) senescence (CSGene, Fridman_up, SenMayo) and (C) fibrosis (Hallmark_EMT, RenalFibrosis) gene sets. (**D)** UMAP of the integrated snRNA-seq, scRNA-seq and snATAC-seq datasets (n = 588,425 cells and nuclei) from the Susztak Lab and KPMP^29^. **(E)** Average expression of NNMT in the injured proximal tubule (iPT) cluster in control and CKD kidney samples from the Susztak Lab and KPMP scRNA-seq data. **(F)** Characteristics of DKD kidney biopsies used for IHC. **(G)** Pearson correlation of eGFR with % IFTA in the DKD kidney biopsies, n=9. **(H)** Characteristics of diabetic and early DKD patients included for urine and serum metabolomics. **(I)** Reaction catalyzed by NNMT. **(J)** LC–HRMS measurement of urinary NAM and MeNAM in diabetic and early DKD patients. **(K)** LC–HRMS measurement of serum NAM, MeNAM and Me6PY in diabetic and early DKD patients. Results shown are mean ± s.e.m., with p-value determined by Student’s two-tailed t-test (J,K).

**SUPPLEMENTARY FIGURE 2.**
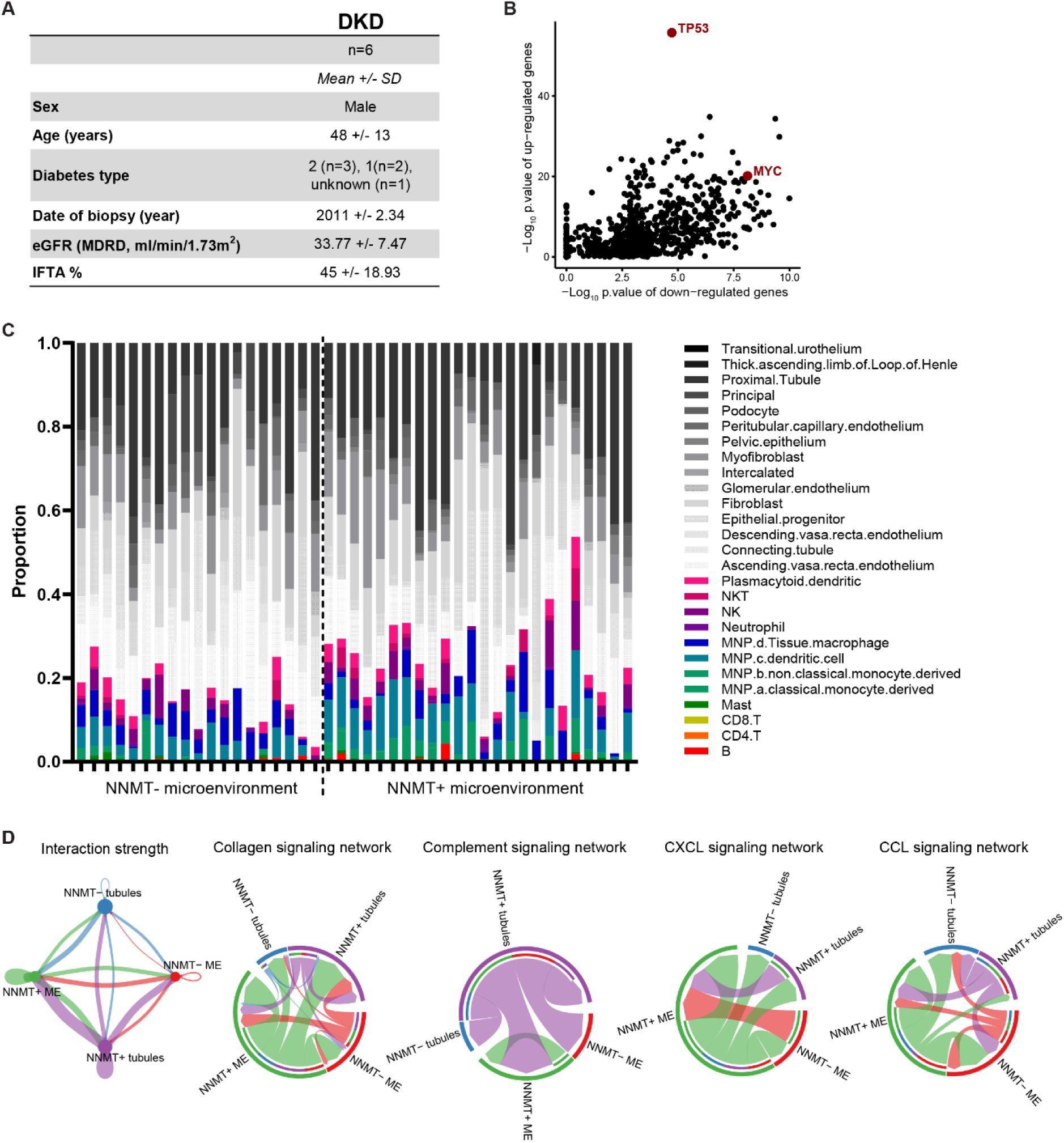
Spatial transcriptomics validates the link between tubular NNMT, senescence and the inflammatory microenvironment. **(A)** Characteristics of DKD kidney biopsies used for spatial transcriptomics. **(B)** Prediction of the transcription factors driving the differentially expressed genes in NNMT+ tubules *vs.* NNMT-tubules using Lisa. **(C)** Cell-type deconvolution analysis of the microenvironment AOIs using public human kidney scRNAseq. **(D)** Cell-cell communication inference analysis between tubule and microenvironment AOIs using CellChat.

**SUPPLEMENTARY FIGURE 3.**
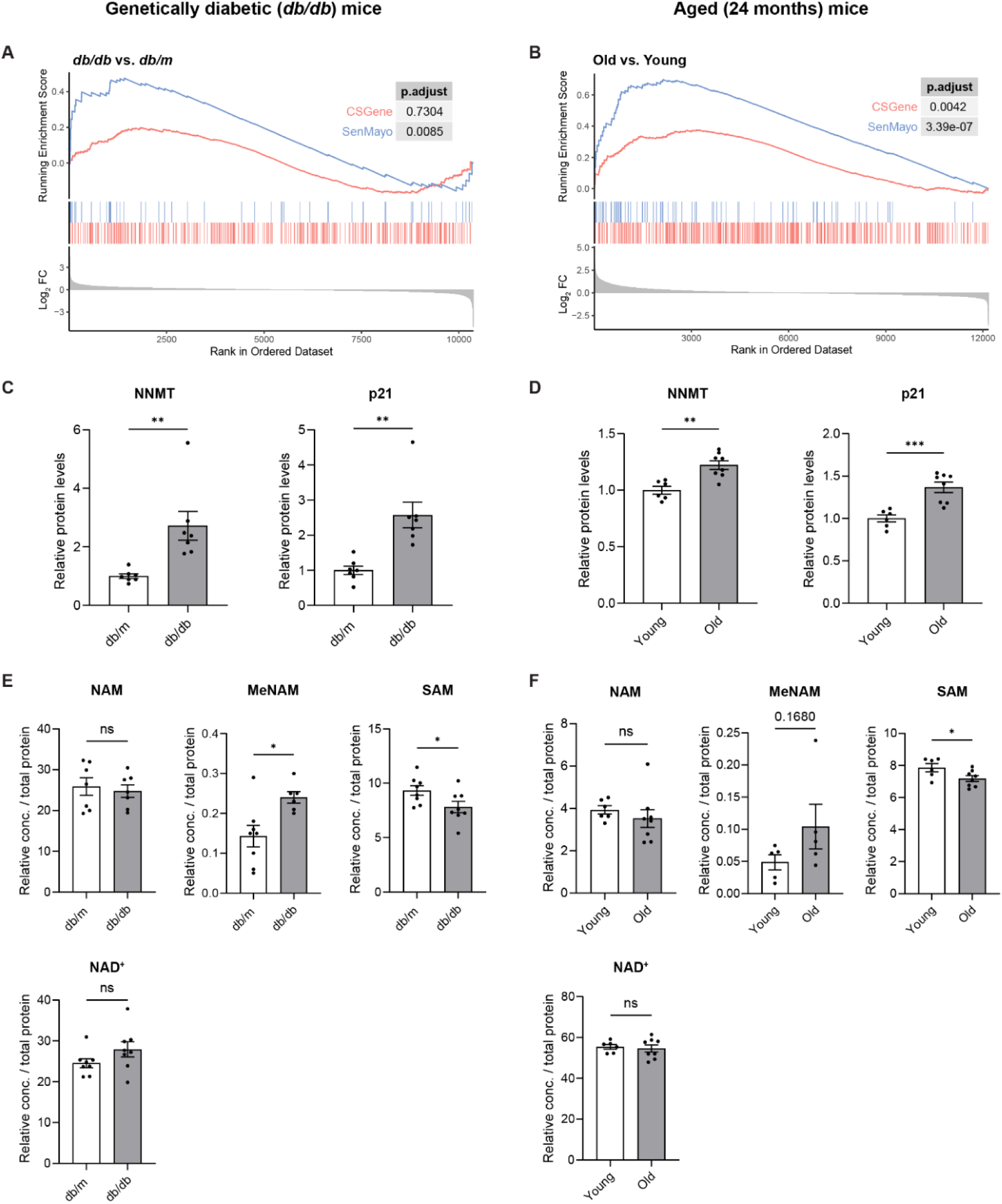
Preclinical models of early CKD show increased renal senescence and NNMT expression. **(A,B)** Senescence GSEA of (A) *db/db vs. db/m* mice kidney and (B) old *vs.* young mice kidney. **(C,D)** Quantification of NNMT and p21 protein levels in (C) *db/m* and *db/db* mice kidney, n=7, and (D) young and old mice kidney, n=6-8. **(E,F)** LC–HRMS measurement of NAM, MeNAM, SAM and NAD^+^ in (E) *db/m* and *db/db* mice kidney, n=6-8, and (F) young and old mice kidney, n=5-8, reanalyzed from the NAM interventional study. Results shown are mean ± s.e.m., with p-value determined by Student’s two-tailed t-test (C-F).

**SUPPLEMENTARY FIGURE 4.**
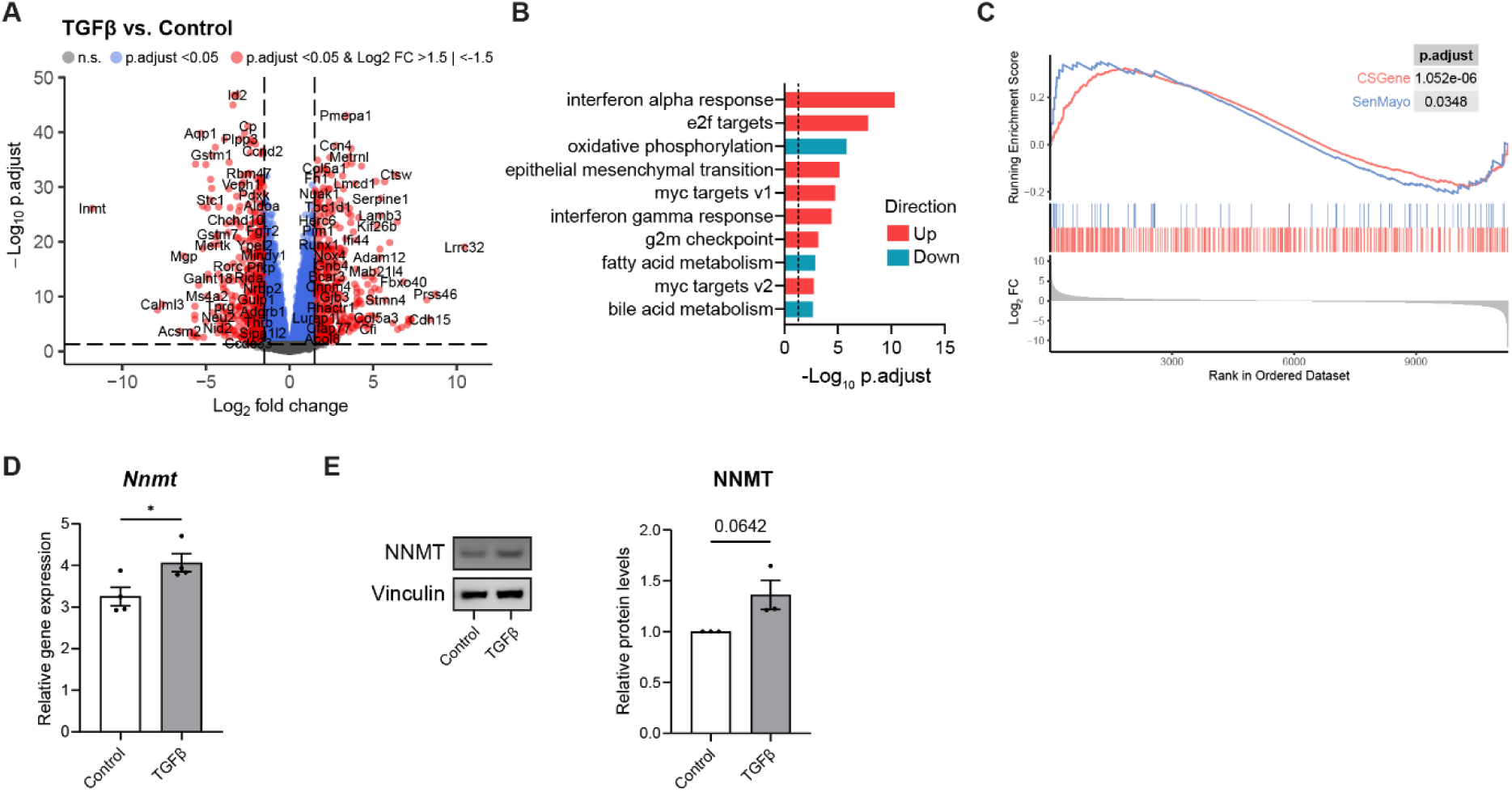
TGFβ stimulation of tubular epithelial cells recapitulates cellular senescence, EMT and increased NNMT expression. (A-C) Transcriptomic analysis of TGFβ-treated tubular epithelial cells *vs.* control cells, n=3: (A) volcano plot of differentially expressed genes, (B) GSEA of the mouse hallmark curated gene set collection from MSigDB, showing top 10 gene sets with the lowest p.adjust, and (C) GSEA of senescence gene sets (CSGene, SenMayo). (D) mRNA expression of *Nnmt* in control cells and cells treated with TGFβ, n=4. (E) Western blot analysis of NNMT and vinculin and quantification of NNMT protein levels in control cells and cells treated with TGFβ, n=3. Results shown are mean ± s.e.m., with p-value determined by Student’s two-tailed t-test (D,E).

**SUPPLEMENTARY FIGURE 5.**
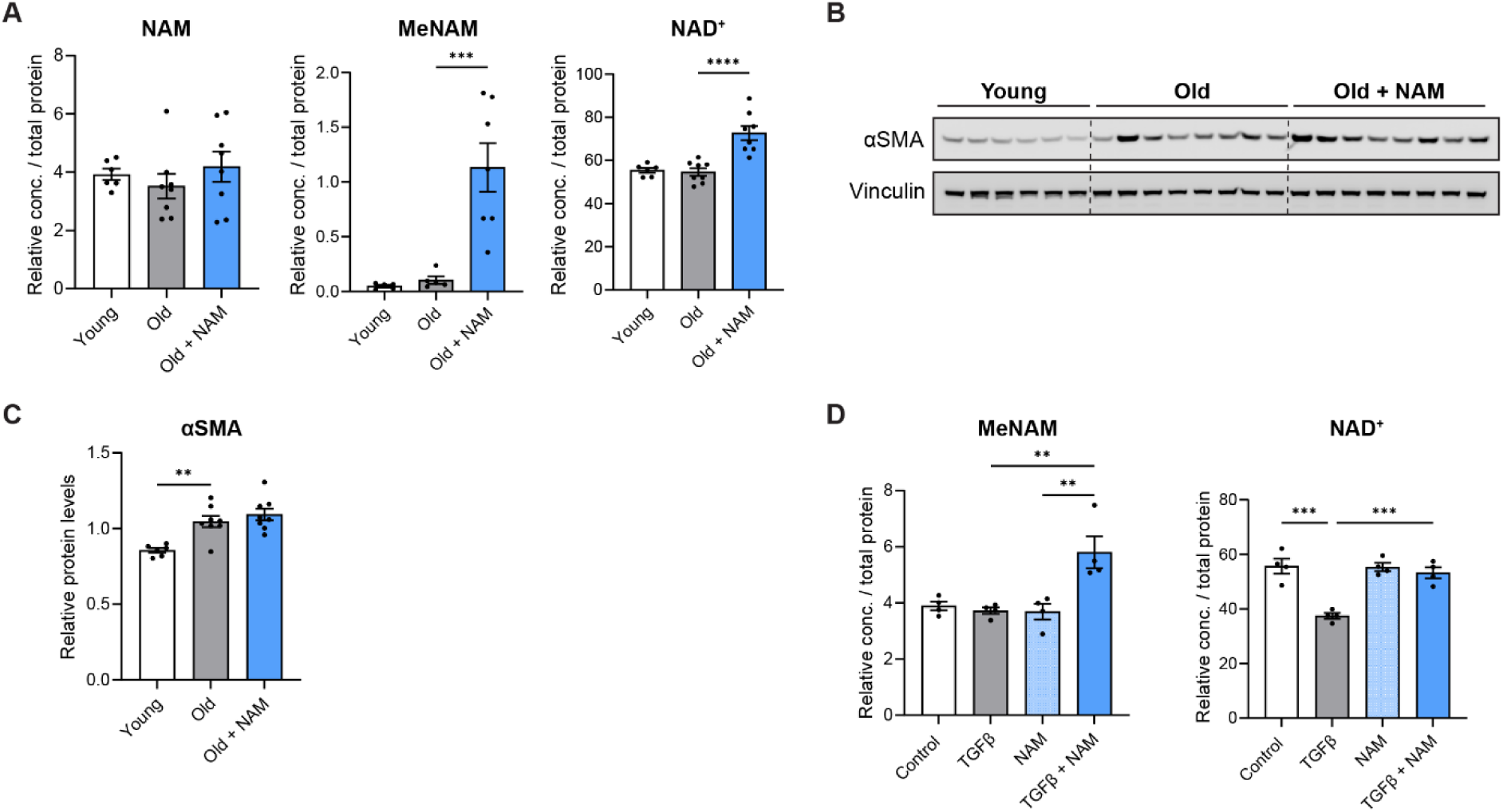
NAM supplementation boosts NAM/NAD^+^ metabolism without preventing fibrosis and senescence. (A) LC–HRMS measurement of NAM, MeNAM and NAD^+^ in the kidney of young mice, aged mice, and aged mice supplemented with dietary NAM for 14 weeks, n=5-8. (B) Western blot analysis of αSMA and vinculin in the kidney of mice as in A and (C) quantification of αSMA protein levels, n=6-8. (D) LC–HRMS measurement of MeNAM and NAD^+^ in tubular epithelial cells after 48-h TGFβ treatment, with or without co-treatment with NAM, n=4. Results shown are mean ± s.e.m., with p-value determined by one-way ANOVA with Šídák’s post-hoc test (A,C,D).

**SUPPLEMENTARY FIGURE 6.**
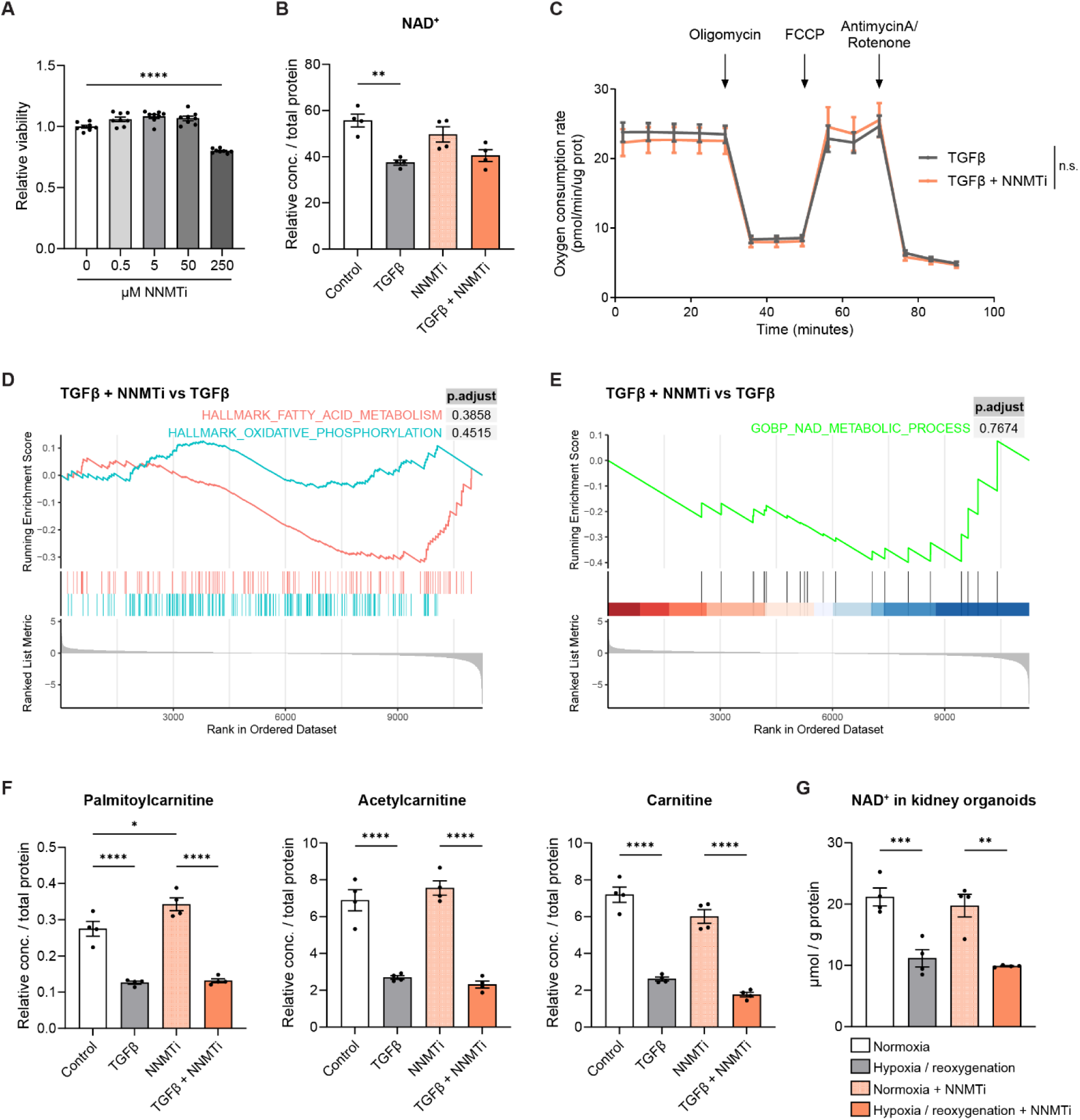
NNMT inhibition does not modulate NAD^+^ levels and mitochondrial metabolism. **(A)** MTT cell viability assay of tubular epithelial cells treated 48 hours with various NNMTi concentrations, n=8. **(B)** LC–HRMS measurement of NAD^+^ in control cells, cells treated with TGFβ, NNMTi, or both, n=4. **(C)** Seahorse-based respiration assay profile from cells stimulated with TGFβ with and without NNMTi, n=6. **(D,E)** GSEA of Log2 fold change ranked genes between TGFβ and NNMTi stimulated cells and TGFβ stimulated cells showing no enrichment in (D) fatty acid metabolism and oxidative phosphorylation and (E) NAD^+^ metabolic process gene sets. **(F)** LC–HRMS measurement of palmitoylcarnitine, acetylcarnitine and carnitine levels in cells as in B, n=4. **(G)** NAD^+^ levels measured in iPSC-derived kidney organoids in normoxia or hypoxia/reoxygenation conditions with or without NNMTi, n=4. Results shown are mean ± s.e.m., with p-value determined by Student’s two-tailed t-test (C) or one-way ANOVA with Šídák’s post-hoc test (A,B,F,G).

